# Harnessing the functional diversity of plant cystatins to design inhibitor variants highly active against herbivorous arthropod digestive proteases

**DOI:** 10.1101/2021.07.22.453419

**Authors:** Jonathan Tremblay, Marie-Claire Goulet, Juan Vorster, Charles Goulet, Dominique Michaud

## Abstract

Protein engineering approaches have been proposed to improve the inhibitory properties of plant cystatins against herbivorous arthropod digestive proteases. These approaches typically involve the site-directed mutagenesis of functionally relevant amino acids, the production and selection of improved inhibitory variants by molecular phage display procedures, or the design of bi/multifunctional translational fusions integrating one or several cystatin inhibitory domains. Here, we propose a new approach where the function-related structural elements of a cystatin are substituted by the corresponding elements of an alternative cystatin. Cys protease inhibitory assays were first performed with 20 representative plant cystatins and model Cys proteases, including herbivorous arthropod digestive proteases, to appreciate the extent of functional variability among plant cystatin protein family members. The most, and less, potent of these cystatins were then used as ‘donors’ of structural elements to create hybrids of tomato cystatin SlCYS8 used as a model ‘recipient’ inhibitor. Our data confirm the wide variety of cystatin protease inhibitory profiles among plant taxa. They also demonstrate the usefulness of these proteins as a pool of discrete structural elements for the design of cystatin variants with improved potency against herbivorous pest digestive Cys proteases.

## INTRODUCTION

Protease inhibitors of the cystatin protein superfamily play various roles in plants, from the regulation of cysteine (Cys) proteases in seeds and senescent organs to the inhibition of exogenous digestive proteases upon arthropod herbivory or pathogenic infection (Arai et al. 2002; Benchabane et al. 2010). Cystatins act as reversible pseudosubstrate inhibitors to hinder the active site of target proteases and block their catalytic action on the peptide bonds of protein substrates (Turk and Bode 1991). The inhibitory function of these proteins relies on two structural elements, a central hairpin loop with the conserved pentapeptide motif Gln–X–Val–X–Gly (where X is any amino acid) and a second hairpin loop in the C-terminal region with a conserved Trp residue, which physically interact with amino acid residues in the active site cleft of the target enzyme. A third structural element is also involved, consisting of a flexible, N-terminal amino acid string presenting a conserved Gly–Gly dipeptide motif (Benchabane et al. 2010). This third element, referred to as the N-terminal trunk, interacts with surface residues on the target enzyme to strongly influence the inhibitory potency and specificity of the cystatin towards different protease isoforms (Rasoolizadeh et al. 2016a).

An increasing body of knowledge about the properties and physiological roles of plant cystatins has triggered the development of various biotechnological applications over the years (Tremblay et al. 2019). Studies have shown the potential of these proteins as ectopic regulators of endogenous Cys proteases to regulate storage protein deposition and degradation in reproductive organs (Hwang et al. 2009, Munger et al. 2015), to restore fertility in Cys protease-induced male sterile plants (Shukla et al. 2016) or to avoid the detrimental action of endogenous Cys proteases on heterologous proteins in plants used as bio-factories for clinically relevant proteins (Sainsbury et al. 2013, Jutras et al. 2016, Grosse-Holz et al. 2018, Jutras et al. 2019). Other studies have shown their potential to implement drought, cold or salt tolerance in different crops (Chen et al. 2014, Quain et al. 2014, Tan et al., 2016, 2017a), associated with the induction of abiotic stress-related genes upon recombinant cystatin expression (Munger et al. 2012, Tan et al. 2017b). Most importantly, numerous studies have described the potential of plant cystatins to protect plants from microbial pathogens, root parasitic nematodes and phytophagous arthropods (reviewed in Schlüter et al. 2010, Lima et al. 2015, Macedo et al. 2015, Martinez et al. 2016). Cystatins inhibit digestive Cys proteases secreted in the extracellular milieu of microbial cells or digestive tract of herbivorous arthropods, to cause amino acid shortage, growth delays and eventual death of the pathogenic or herbivorous enemy (Broadway 2001, Lima et al. 2015).

From a physiological standpoint, the actual ability of a cystatin to protect the plant from herbivory is determined by its relative abundance compared to Cys proteases in the target herbivore midgut, by its inhibitory range towards these enzymes, and by any compensatory response induced in the herbivore after ingestion (Rasoolizadeh et al. 2016b). Herbivorous insects have developed effective strategies to avoid the negative effects of dietary protease inhibitors, including the secretion of digestive proteases from different functional classes, the overexpression of proteases following inhibitor uptake, and the production of protease isoforms weakly sensitive to inhibition (Zhu-Salzman and Zeng 2015). A well-documented example is the coleopteran insect pest Colorado potato beetle (*Leptinotarsa decemlineata*), which uses an array of positively selected digestive Cys protease isoforms to process leaf proteins (Vorster et al. 2015). Divergent, if not contradictory, effects have been reported for transgenic potato lines engineered to express cystatins, ranging from major developmental delays and mortality (Lecardonnel et al. 1999, Cingel et al. 2015, Rasoolizadeh et al. 2016b) to compensatory growth and hypertrophic behavior sustained by Cys protease overexpression (Cloutier et al. 1999, 2000, Cingel al. 2015). Possible explanations for such discrepancy among studies include differential expression levels of the recombinant cystatin in leaf tissue, varying stability of this protein in different potato cultivars, distinct inhibitory ranges towards the insect Cys proteases and experimental biases influencing insect fitness. Together, these observations stress the need for a better understanding of complex interactions between wound-inducible cystatins and Cys proteases in plant–insect systems. They also underline the relevance of rational strategies for the molecular improvement of recombinant cystatin inhibitory profiles towards Cys proteases.

Three main approaches are generally adopted for the molecular improvement of plant cystatins (Sainsbury et al. 2012a, van Wyk et al. 2016, Tremblay et al. 2019). The first approach involves site-directed substitution at functionally relevant amino acid sites, the second approach the generation of improved inhibitory variants using phage display or DNA shuffling artificial evolution procedures, and the third approach the design of bi- or multifunctional translational fusions integrating one or more cystatin inhibitory domains. In this study, we explored the potential of a fourth approach based on the substitution of one or more function-related structural elements (SE’s) of the cystatin by the corresponding element(s) of an alternative cystatin. Our goal was to assess the usefulness of potent cystatins from different plant taxa as SE “donors” to generate functional variability among the structural hybrids of a “recipient” cystatin. The idea was to translate the concept of ‘loop replacement design’ (LRD), as described for the engineering of multimeric mammalian antibodies (Clark et al., 2009), to the improvement of single-domain cystatins. Amino acid substitutions in the N-terminal trunk or the inhibitory loops of plant cystatins have proved useful to enhance the inhibitory potency or change the affinity profile of these proteins towards insect or nematode Cys proteases (Urwin et al. 1995, Kiggundu et al. 2006, Goulet et al. 2008, Rasoolizadeh et al. 2016a). We hypothesized that an LRD-like scheme by which the function-related elements of a protein are changed for the corresponding elements of a related protein would represent a welcome complement to site-directed mutagenesis as it would allow conformational changes in the cystatin on a length scale beyond that accessible to single mutations (Clark et al. 2009).

Cystatins are well suited to protein engineering and polypeptide grafting, as illustrated by their stability in fusion with different protein partners (Jutras et al. 2018, Tremblay et al. 2019), the structural stability of model cystatin tomato SlCYS8 bearing a poly-His tag for protein purification in a non-inhibitory loop of the protein scaffold (Sainsbury et al. 2016), the ability of SlCYS8 to stabilize a human protein translational fusion partner *in planta* (Sainsbury et al. 2013), the structural assessment of natural cystatins as a guide for *de novo* protein design (Marcos et al. 2017), and the use of a consensus plant cystatin scaffold to design Affimer binding proteins for a variety of imaging, diagnostic and therapeutic purposes (Tiede et al. 2014, Kyle 2018). Here, we confirm the usefulness of plant cystatins as a reservoir of discrete structural elements for cystatin engineering, and the potential of SE substitutions to create cystatins with improved inhibitory potency against arthropod Cys proteases.

## RESULTS

### Variable contributions of the N-terminal trunk and two inhibitory loops to the protease binding strength of plant cystatins

Docking simulations were performed *in silico* with three protease models to gain some preliminary insight about the functional variability of plant cystatins and the relative contributions of their function-related SE’s to the enzyme–inhibitor complex. Five plant cystatins and the three model Cys proteases papain, human cathepsin L and Colorado potato beetle intestain D4 (IntD4) (Vorster et al. 2015) were selected for the simulations, for a total of 15 protease–cystatin complexes and 45 protease–SE interactions (**Table 1**). Structure models were first built for the proteases and the cystatins by homology modelling with the solved structures of human cathepsin L (Ljunggren et al. 2007) and oryzacystatin I (OsCYS1) (Nagata et al. 2000), respectively. Protease–cystatin interactions were then simulated using the Z-Dock algorithm of Chen et al. (2003), by homology to the solved structure of papain in complex with human stefin B (Protein Data Bank Accession No. 1STF). In line with variable sequences in the functional regions of both the cystatins and the proteases, amino acid residues predicted to contribute to the binding process differed from one cystatin to another for a given protease, and from one protease to another for a given cystatin (**Supplementary Figure 1**). Accordingly, total binding energies differed for the 15 protease–cystatin complexes, from an inferred total energy value of –544 kcal/mol for maize cystatin ZmCYS1 interacting with papain to an energy value of –1234 kcal/mol indicating a stronger interaction between the same cystatin and cathepsin L (**Table 1**, **Supplementary Table 1**).

**Table 1.** Binding energies inferred *in silico* for Cys proteases papain, human cathepsin L and *L. decemlineata* Intestain D4 interacting with different plant cystatins ^1^

| Cystatin / Protease | Interaction energy (kcal/mol) |  |  |  |
| --- | --- | --- | --- | --- |
|  | N-ter trunk | Loop 1 | Loop 2 | Total |
| **SICYS8** |  |  |  |  |
| . Papain | −141.0 | −105.9 | −385.6 | −632.5 |
| . Cathepsin L | −386.6 | −133.5 | −408.8 | −928.9 |
| . Intestain D4 | −333.2 | −84.4 | −240.1 | −657.7 |
| **SICYS9** |  |  |  |  |
| . Papain | −133.6 | −180.1 | −401.6 | −715.3 |
| . Cathepsin L | −389.0 | −327.3 | −274.5 | −990.8 |
| . Intestain D4 | −302.2 | −92.1 | −271.7 | −666.0 |
| **OsCYS1** |  |  |  |  |
| . Papain | −151.1 | −170.0 | −317.9 | −639.0 |
| . Cathepsin L | −419.7 | −273.3 | −290.8 | −983.8 |
| . Intestain D4 | −341.1 | −104.0 | −289.1 | −734.2 |
| **GmCYS2** |  |  |  |  |
| . Papain | −333.8 | −188.7 | −483.5 | −1006.0 |
| . Cathepsin L | −412.8 | −303.6 | −218.0 | −934.4 |
| . Intestain D4 | −346.6 | −165.8 | −192.6 | −705.0 |
| **ZmCYS1** |  |  |  |  |
| . Papain | −131.9 | −136.4 | −276.0 | −544.3 |
| . Cathepsin L | −591.2 | −322.4 | −320.4 | −1234.0 |
| . Intestain D4 | −326.4 | −76.8 | −202.7 | −605.9 |

<sup>1</sup> Data are the sum of binding energy values inferred for the complement of protease–cystatin interacting residues associated with the N-terminal trunk (N-ter), the first inhibitory loop (Loop 1), the second inhibitory loop (Loop 2) or the whole cystatin (Total) (See **Supplementary Table 1** for binding energy values at the amino acid level). Cystatin interacting residues are identified in **Supplementary Figure 1** for the three model proteases.

In support to previously described models indicating variable contributions of the N-terminal trunk and two inhibitory loops to the protease binding process (Vorster et al. 2010), binding energies assigned to the three structural elements differed depending on the cystatin or the protease considered (**Table 1**). For instance, a binding energy value of –334 kcal/mol accounting for 33% of the complex total binding energy was inferred for the N-terminal trunk of soybean cystatin GmCYS2 interacting with papain, compared to weaker energy values and relative contributions of less than 25% for the N-terminal trunks of tomato cystatins SlCYS8 and SlCYS9, OsCYS1 and ZmCYS1 interacting with the same enzyme (**Figure 1**). Likewise, an energy value of –402 kcal/mol accounting for 56% of the total was calculated for the second inhibitory loop of SlCYS9 interacting with papain, compared to weaker binding energies (and smaller relative contributions) of –275 kcal/mol (28%) and –272 kcal/mol (41%) for the same cystatin interacting with cathepsin L and IntD4, respectively (**Table 1**).

**Figure 1.**
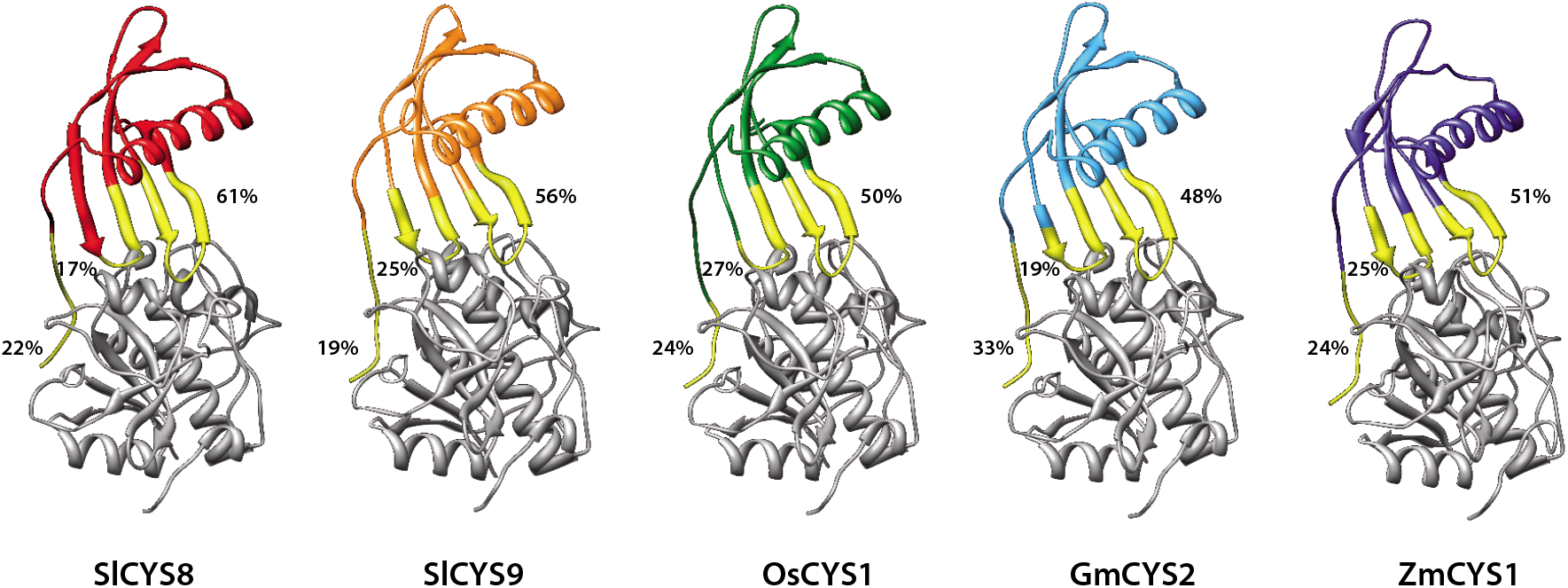
Docking models for tomato SlCYS8 and SlCYS9, rice OsCYS1 (oryzacystatin I), soybean GmCSY2 and corn ZmCYS1 interacting with papain (in grey). Cystatin residues physically interacting with the target enzyme are highlighted in yellow. Numbers indicate the relative contributions of the N-terminal trunk, first inhibitory loop and second inhibitory loop to the binding process, as inferred from Table 1 (total = 100%).

### Functional variability among plant cystatin protein family members

Protease inhibitory assays were conducted with cystatins of different plant taxa to empirically support our *in silico* assumptions suggesting functional variability among plant cystatins, to confirm the potential of these proteins as a source of SE’s for cystatin improvement, and to identify potent cystatin donors for the SE substitution experiments. A multiple sequence alignment was generated with 262 cystatin primary sequences available in the National Center for Biotechnology Information (NCBI) database, using tomato SlCYS8 as a reference (Goulet et al. 2008). Double-stranded DNA fragments, or ‘g-blocks’, were then produced for 30 of the cystatins, chosen based on their distribution in different branches of the resulting phylogenetic tree (**Supplementary Figure 2**) and their belonging to different subgroups of the plant cystatin family (Benchabane et al. 2010). The DNA fragments were used as coding gene templates for heterologous expression in *E. coli* and affinity purification using the GST gene fusion (Sainsbury et al. 2016). A total of 20 cystatins or cystatin domains deemed representative of the plant cystatin protein family (**Supplementary Figure 2**) were recovered under a stable form and used as test inhibitors for the protease assays (**Table 2**).

**Table 2.**
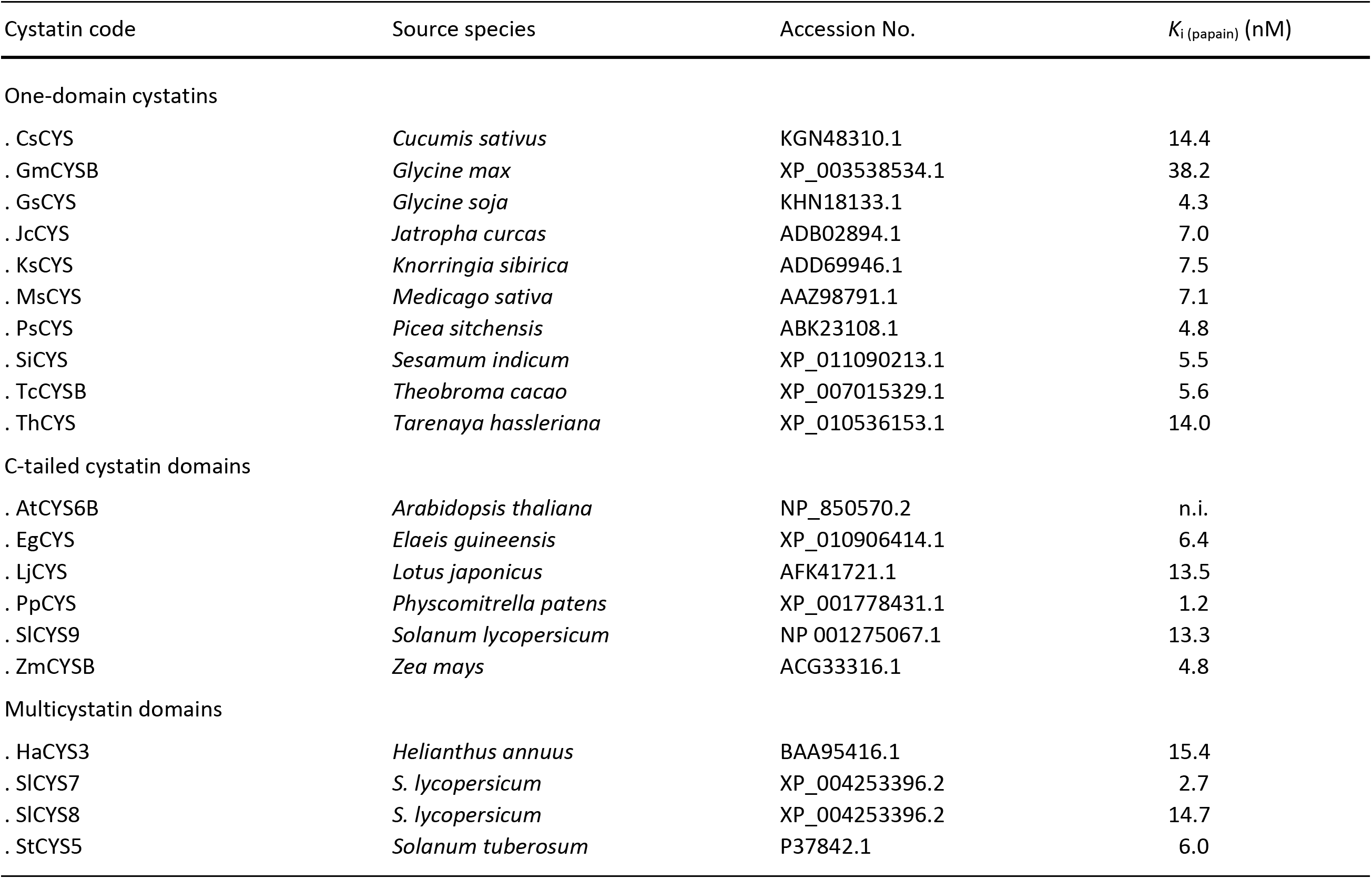
Plant cystatins selected for the functional studies and their calculated *K*_i_ values for model Cys protease papain

In agreement with our *in silico* models indicating variable binding energies for SlCYS8 and other plant cystatins interacting with papain (**Table 1**), inhibition constant (*K*_i_) values against papain (*K*_i (papain)_) differed by more than one order of magnitude from one cystatin to another, from 1.2 nM for *Physcomitrella patens* C-tailed cystatin domain PpCYS or 2.7 nM for tomato multicystatin domain SlCYS7, to 38.2 nM for soybean GmCYSB or even no measurable inhibitory activity for Arabidopsis AtCYS6B (**Table 2**). Similarly, the 20 cystatins showed variable inhibitory potency against midgut cathepsin L-like (*Z*-Phe–Arg-MCA hydrolyzing) Cys proteases of *L. decemlineata* and the acarian herbivore generalist two-spotted spider mite, *Tetranychus urticae* (**Figure 2**). For instance, *P. patens* cystatin PpCYS and potato multicystatin domain StCYS5 showed strong inhibitory activity against these proteases at low (20 nM) concentration, in sharp contrast with cucumber cystatin CsCYS and barley cystatin HaCYS3 showing negligible activity. Not surprisingly given the high specificity of Cys protease–cystatin interactions at the submolecular level, several cystatins showed variable effects depending on the protease tested (**Table 2**, **Figure 2**). This was observed for instance with AtCYS6B showing no activity against papain and *L. decemlineata* cathepsin L-like enzymes but easily measurable activity against *T. urticae* cathepsin L-like enzymes, or with *Glycine soja* GsCYS efficiently inhibiting papain and *T. urticae* proteases but showing weaker activity against the *L. decemlineata* enzymes.

**Figure 2.**
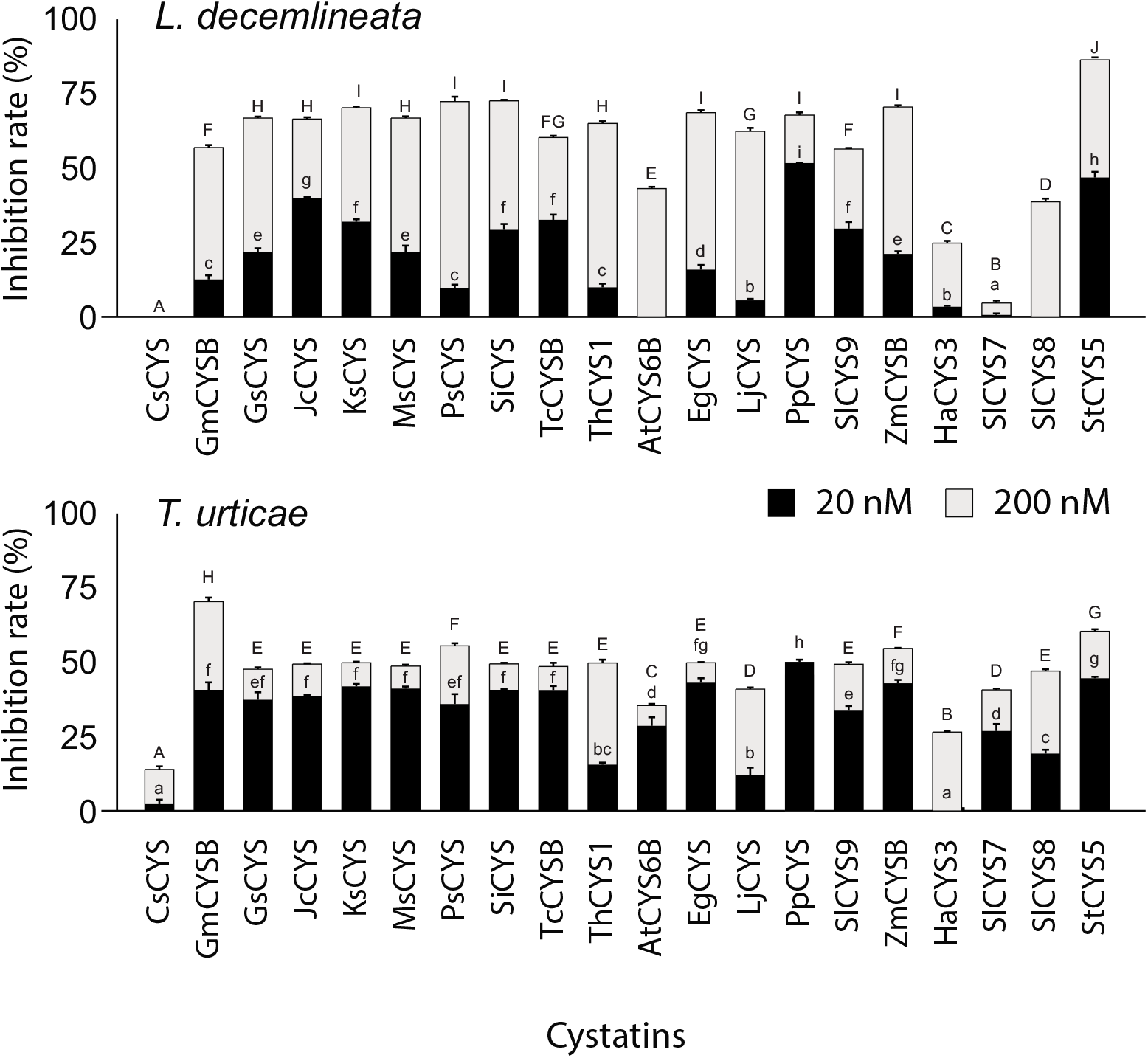
Inhibition of *L. decemlineata* and *T. urticae* of *Z*-Phe–Arg-MCA-hydrolyzing (cathepsin L-like) enzymes by 20 representative members of the plant cystatin protein family (as identified in **Table 2**). Data are expressed as relative inhibitory rates compared to the inhibitory rate measured with E-64 (100%), a broad-spectrum diagnostic inhibitor for Cys proteases of the C1A (papain) family. Inhibitory assays were conducted with limiting (20 nM) or excess (200 nM) concentrations of cystatin. Each bar is the mean of three independent (biological) replicates ± SE.

### A generic scheme for plant cystatin SE substitutions

An SE substitution strategy was designed to determine whether the variable inhibitory effects of plant cystatins against Cys proteases could be formally associated with the primary structures of their N-terminal trunk and inhibitory loops as suggested by our docking inferences, and whether plant cystatins as a group would represent a useful pool of discrete structural elements for the design of cystatin variants with improved potency against herbivorous pest digestive Cys proteases. Tomato SlCYS8 was used as a recipient protein model for hybrid design given its reported suitability for protein engineering (Goulet et al. 2008, Sainsbury et al. 2013, 2016, Jutras et al. 2018) and moderate activity against Cys proteases compared to other plant cystatins (**Table 2** and **Figure 2**). *P. patens* PpCYS and potato StCYS5 were used as donors given their strong inhibitory potency against *L. decemlineata* proteases, and hence the expected potential of their function-related structural elements for SlCYS8 improvement. Cucumber CsCYS, weakly active against the arthropod cathepsin L-like enzymes (**Figure 2**), was selected as a ‘flawed’, negative control donor to further confirm the potential of inherently efficient inhibitors such as PpCYS and StCYS5 as loop donors to generate potent cystatins.

The SE hybrids were designed *in silico* by substituting the sequence(s) of SlCYS8 N-terminal trunk, first inhibitory loop (L1) and/or second inhibitory loop (L2) by the corresponding element(s) of PpCYS, StcYS5 or CsCYS (**Figure 3**). The N- and C-terminal boundaries of each structural element were defined based on their distance relative to conserved amino acid motifs essential for activity in the transferred element, in such a way as to also include all amino acids assumed to physically interact with amino acid residues of the target enzyme (Vorster et al. 2015) (**Figure 3**). More specifically, the N-terminal trunks were devised based on the Gly–Gly (–GG–) motif characteristic of the N-terminal region of plant cystatins, the first inhibitory loops based on the conserved pentapeptide motif Gln–X– Val–X–Gly (–QxVxG–) interacting with specific residues in the active site of the target protease, and the second inhibitory loops based on the conserved Trp residue also interacting with specific residues in the active site cleft (Benchabane et al. 2010). Twenty-one cystatin variants were designed overall, including all seven structural element combinations possible for each of the three cystatin donors (**Table 3** and **Supplementary Table 2**). DNA g-blocks were produced for the 21 hybrids and used as templates for bacterial expression and affinity purification using the GST gene fusion. As for the original cystatins above, some hybrids could not be properly expressed under our experimental conditions, likely due to deficient stability in a foreign cellular environment during heterologous expression. Overall, 16 hybrids were produced in a form suitable for protease inhibitory assays with papain and the arthropod proteases (**Table 3**), a large enough number of variants to draw conclusive trends about the potential of SE substitutions for cystatin engineering.

**Table 3.** SlCYS8 hybrids designed for the functional assays using the N-terminal trunk and/or inhibitory loops of potato cystatin domain StCYS5, *P. patens* cystatin PpCYS or cucumber cystatin CsCYS

| SICYS8 hybrid | Transferred structural elements (•) |  |  |
| --- | --- | --- | --- |
|  | N-ter trunk | Loop1 | Loop 2 |
| . StCYS5 elements |  |  |  |
| S-N <sup>1</sup> | • |  |  |
| S-1 |  | • |  |
| S-2 |  |  | • |
| S-N1 | • | • |  |
| S-N2 | • |  | • |
| S-12 |  | • | • |
| S-N12 <sup>2</sup> | • | • | • |
| . PpCYS elements |  |  |  |
| P-N | • |  |  |
| P-1 |  | • |  |
| P-2 |  |  | • |
| P-N1 | • | • |  |
| P-N2 | • |  | • |
| P-12 |  | • | • |
| P-N12 <sup>2</sup> | • | • | • |
| . CsCYS elements |  |  |  |
| C-N | • |  |  |
| C-1 <sup>2</sup> |  | • |  |
| C-2 |  |  | • |
| C-N1 <sup>2</sup> | • | • |  |
| C-N2 | • |  | • |
| C-12 <sup>2</sup> |  | • | • |
| C-N12 | • | • | • |

<sup>1</sup> S, P and C stand for StCYS5, PpCYS and CsCYS, respectively; N, 1 and 2 for the N-terminal trunk, first inhibitory loop and second inhibitory loop of the donor cystatin.
<sup>2</sup> Unsuccessfully produced in *E. coli* and not considered further for the functional assays.

**Figure 3.**
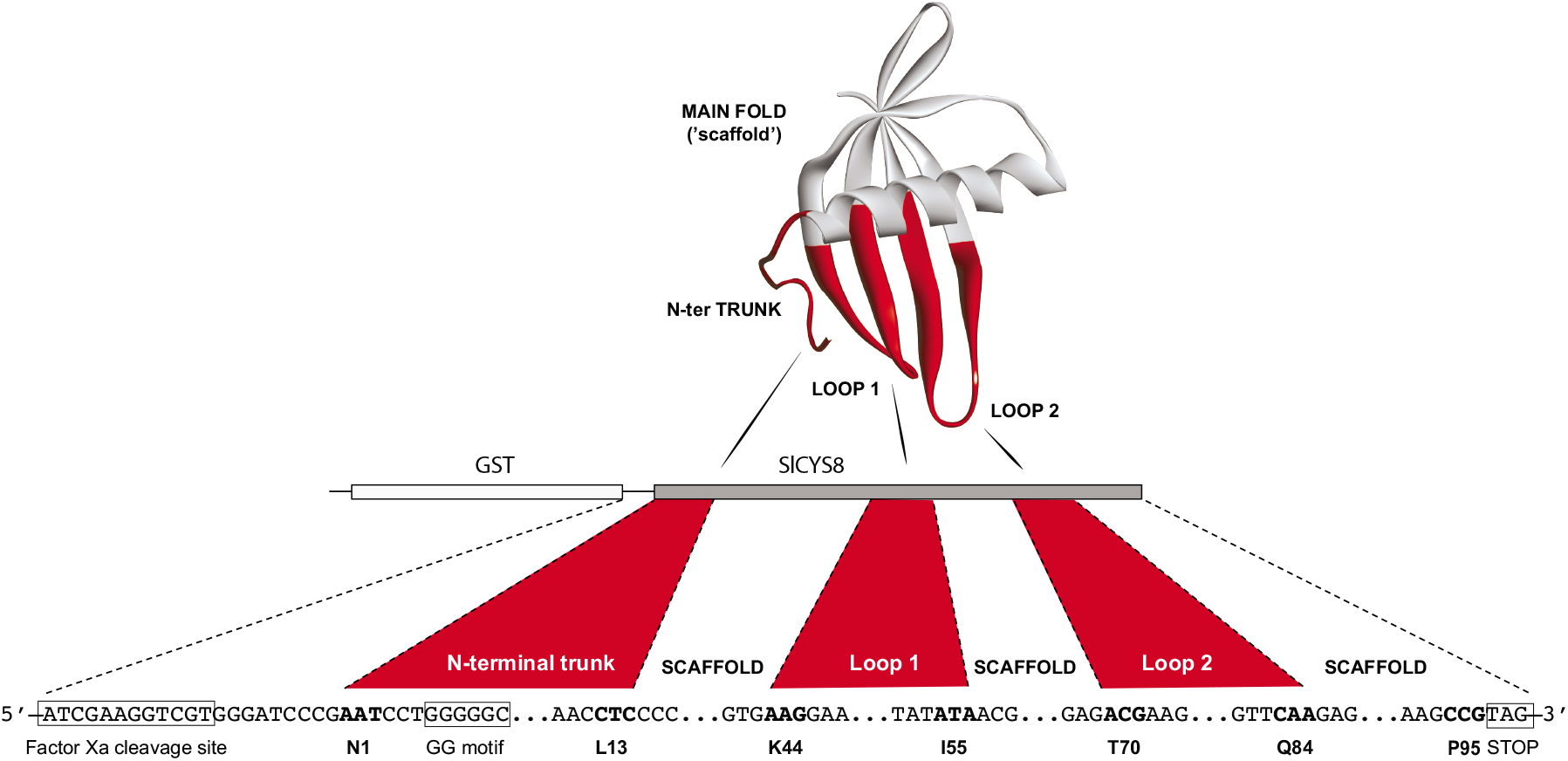
A generic scheme for plant cystatin SE substitutions. The cystatin hybrids were first designed *in silico* by substituting the N-terminal trunk, first inhibitory loop [Loop 1] and/or second inhibitory loop [Loop 2] (in red) of tomato SlCYS8 used as a recipient (or scaffold) by the corresponding element(s) of potato StCYS5, *P. patens* PpCYS or cucumber CsCYS used as donors. DNA g-blocks synthesized for the resulting hybrids were then inserted in a modified pGEX-3X vector, downstream of a GST tag coding sequence, for heterologous expression in *E. coli* and affinity-purification. Numbered amino acids under the gene sequence correspond to the N- and C-terminal amino acids of the three substituted elements. The structural model for SlCYS8 was built using the Discovery Studio 2.5 protein modelling software (Accelrys, San Diego CA) based on the spatial coordinates of rice OsCYS1 (Protein Data Bank Accession 1EQK) (Nagata et al. 2000).

### SE substitutions for the molecular improvement of tomato SlCYS8

Papain inhibitory assays were conducted to measure the impact of N-terminal trunk and inhibitory loop substitutions on the inhibitory activity of SlCYS8. In line with the variable efficiencies of SlCYS8 and donor cystatins against papain, *K*_i_ values for this enzyme differed from one hybrid to another (**Figure 4**). The most potent hybrids were hybrids P-1 and P-N1, both including the first loop of PpCYS, that showed *K*_i_ values for papain almost seven times smaller than the *K*_i_ value determined for SlCYS8. The less potent hybrid was hybrid C-N12, with the three functional elements of CsCYS, that showed no measurable activity against papain. Overall, most substitutions involving the structural elements of StCYS5 and PpCYS, both more potent than SlCYS8 and CsCYS against papain (see **Table 2**), showed decreased *K*_i_ values for this enzyme compared to the original inhibitor, unlike substitutions with the structural elements of CsCYS giving a more contrasted picture.

**Figure 4.**
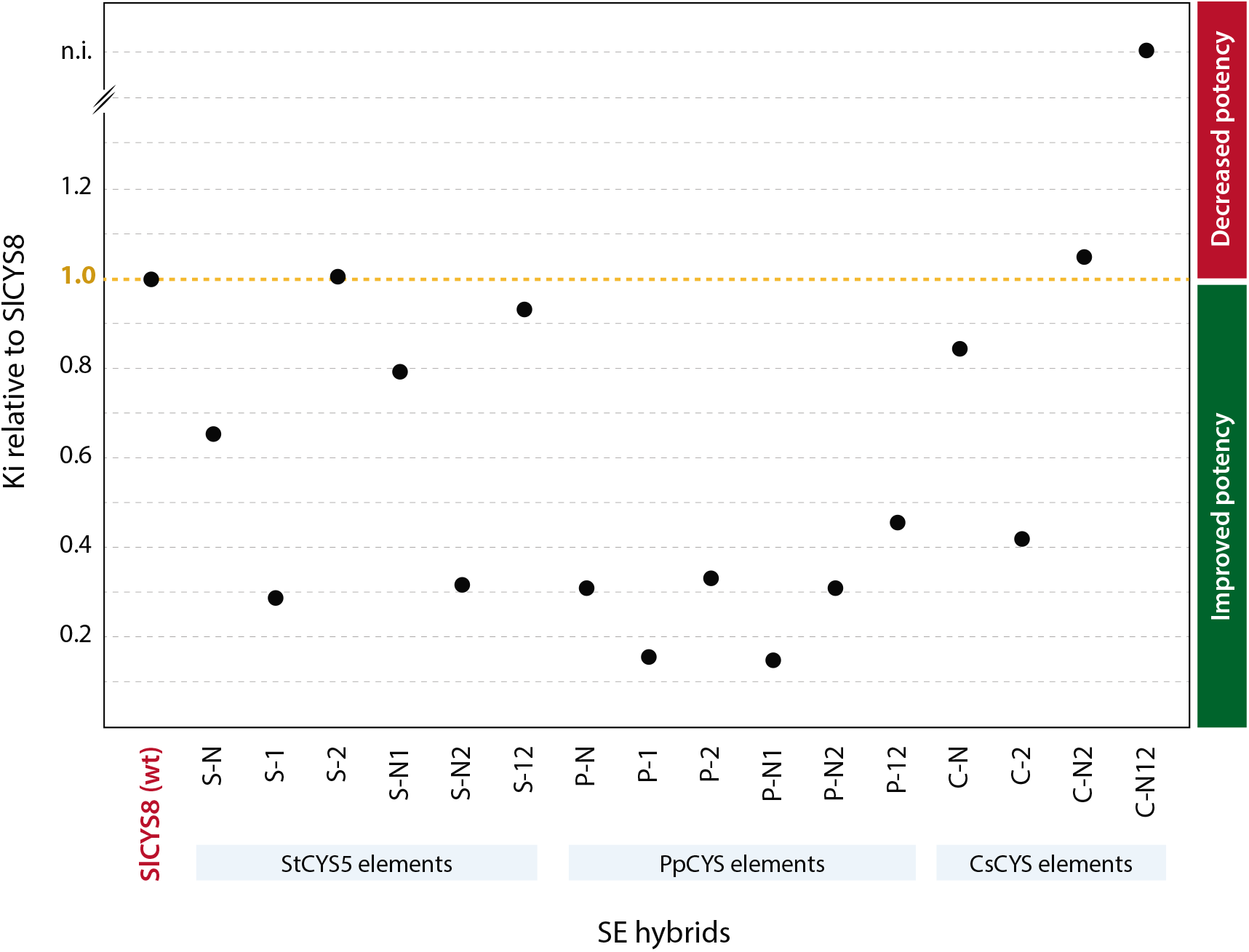
*K*_i (papain)_ values for the SlCYS8 SE hybrids, relative to the *K*_i (papain)_ value for wild-type SlCYS8. A ratio greater than 1.0 indicates a negative impact, and a ratio lower than 1.0 a positive impact, of the element substitution(s) on papain inhibitory activity. S, P and C stand for StCYS5, PpCYS and CsCYS, respectively; N, 1 and 2 for the N-terminal trunk, first inhibitory loop and second inhibitory loop of the donor cystatin.

Similar trends were observed with the two arthropod proteases (**Figure 5** and **Supplementary Figure 3**). Most substitutions for the structural elements of StCYS5 and PpCYS, two potent inhibitors of *L. decemlineata* cathepsin L-like enzymes, strongly improved the inhibitory potency of SlCYS8 against these proteases, unlike substitutions for structural elements of CsCYS having a general negative impact (**Figure 5**, upper panel). This was illustrated for instance by an anti-cathepsin L activity of SlCYS8 increased by ten times following the substitution of its two inhibitory loops by the two loops of StCYS5 (Hybrid S-12) or those of PpCYS (Hybrid P-12), in sharp contrast with the systematic low inhibitory potency of hybrids bearing the N-terminal trunk and/or inhibitory loop(s) of cucumber CsCYS (Hybrids C-N, C-2, C-N2 and C-N12). Changing the structural elements of SlCYS8 by those of StCYS5 or PpCYS had little impact overall for the acarian proteases (**Supplementary Figure 3**), likely explained by the roughly similar inhibitory efficiencies of SlCYS8, StCYS5 and PpCYS against *T. urticae* cathepsin L-like enzymes (see **Figure 2**). By comparison, substitutions for the structural elements of CsCYS had a general negative impact, in accordance with the weak activity of this cystatin against *T. urticae* cathepsin L enzymes compared to SlCYS8 and the other two donor cystatins. Most interestingly, grafting the N-terminal trunk and first inhibitory loop of SlCYS5 to SlCYS8 (Hybrid S-N1) increased its inhibitory potency by more than 20 times against *L. decemlineata* cathepsin B-like enzymes (**Figure 5**, lower panel). This improved inhibitory rate was more than three times the inhibitory rate observed for StCYS5 used at low concentration, suggesting the potential of SE substitutions not only to improve the inhibitory potency of a cystatin against its natural protease targets but also to broaden its inhibitory range to other Cys proteases.

**Figure 5.**
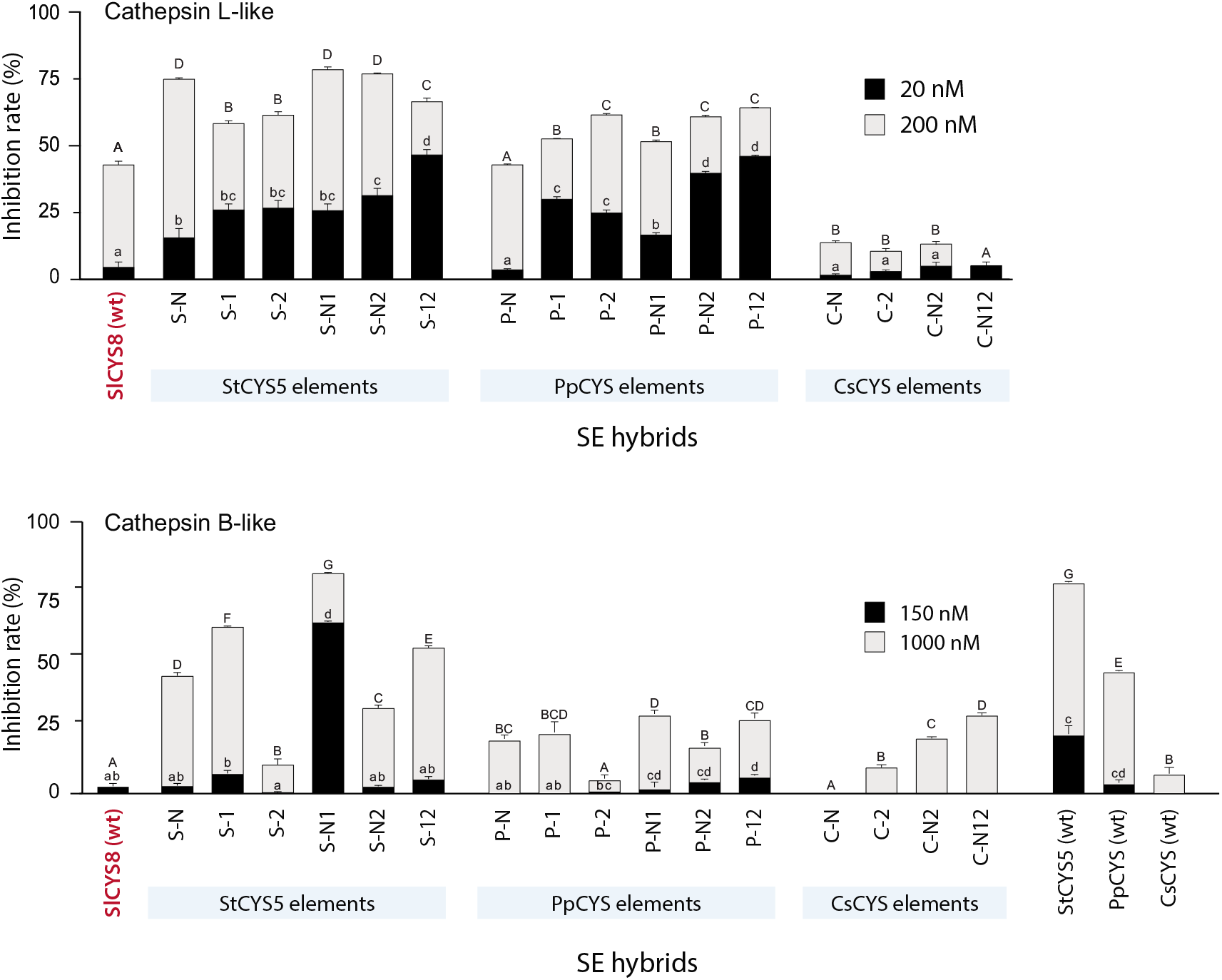
Inhibition of *L. decemlineata Z*-Phe–Arg-MCA-hydrolyzing (cathepsin L-like) and *Z*-Phe–Arg-MCA-hydrolyzing (cathepsin B-like) enzymes by the SlCYS8 SE hybrids. Data are expressed as relative inhibitory rates compared to the inhibitory rate measured with E-64 (100%). Inhibitory assays were conducted with limiting (20 nM [cathepsin L], 150 nM [cathepsin B activity]) or excess (200 nM [cathepsin L], 1 μM [cathepsin B]) concentrations of cystatin. Each bar is the mean of three independent (biological) replicates ± SE. Different letters (lower case for the limiting concentrations, capital for the excess concentrations) indicate significantly different inhibitory rates among cystatin hybrids (post-ANOVA Tuckey’s tests, with an alpha value of 5%). S, P and C stand for StCYS5, PpCYS and CsCYS, respectively; N, 1 and 2 for the N-terminal trunk, first inhibitory loop and second inhibitory loop of the donor cystatin.

*K*_i (papain)_ distribution maps were drawn to compare the overall impact of our SE substitutions strategy on SlCYS8 inhibitory activities with the impact of our site-directed mutagenesis approach involving single substitutions at functionally relevant, positively selected amino acid sites (Kiggundu et al. 2006) (**Figure 6**). An overall coefficient of variation (CV) of 53% was calculated for the relative *K*_i (papain)_ values of a previously described collection of 24 SlCYS8 single mutants bearing an alternative amino acid at positively selected sites Pro-2 (P2) or Thr-6 (T6) in the N-terminal trunk (Goulet et al. 2008). By comparison, CV values of 104% and 73% were calculated for the here tested 20 original (natural) cystatins and 16 SE hybrids, respectively. More specifically, *K*_i (papain)_ values for the single mutants were improved by 31% overall relative to wild-type SlCYS8, smaller than the average improvement rate of 154% observed for the SE hybrids (post-ANOVA Fisher’s LSD test, *P*=0.02).

**Figure 6.**
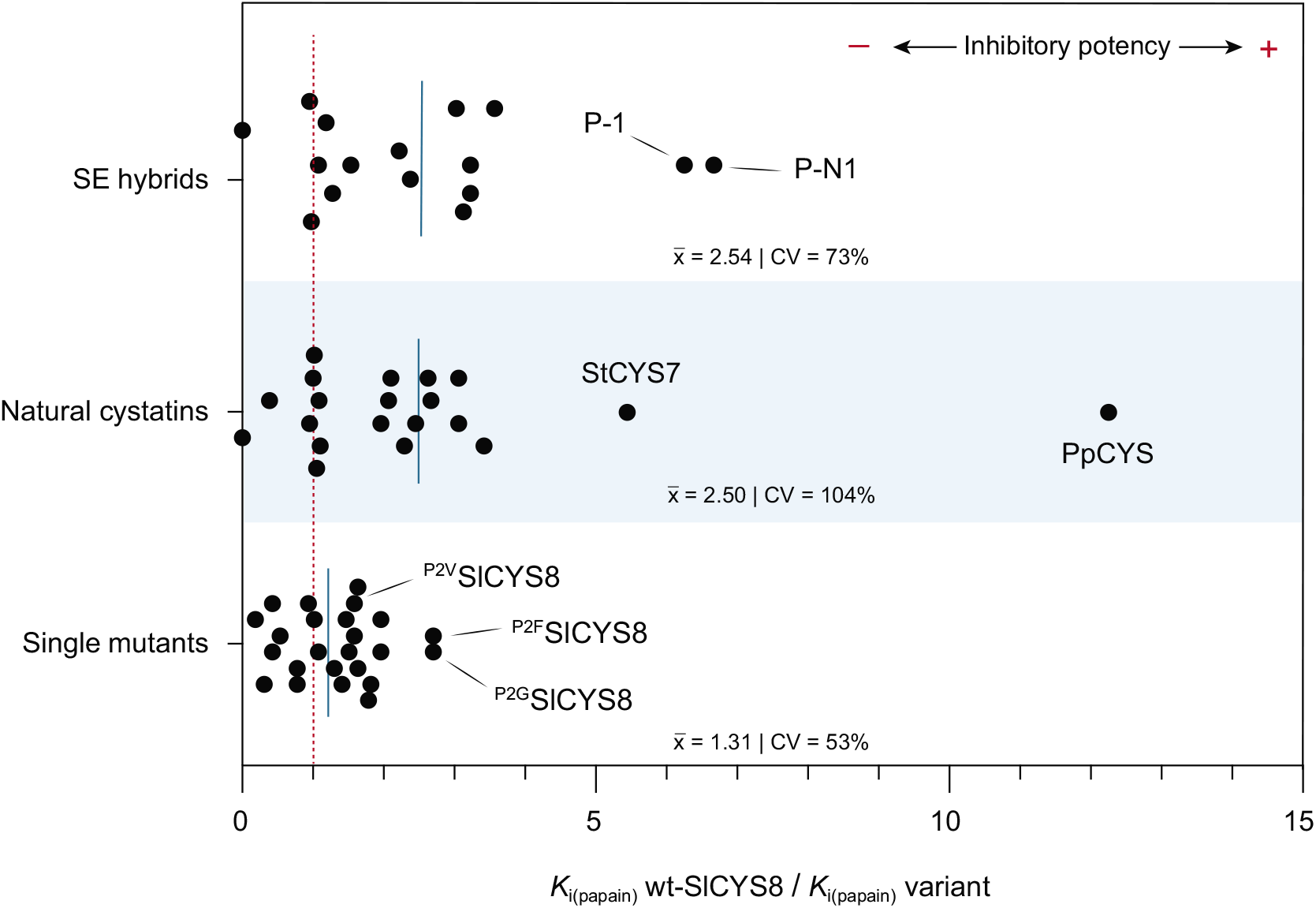
Functional variability among populations of SlCYS8 variants produced by site-directed mutagenesis at positively selected amino acid sites (Goulet et al. 2008), SlCYS8 SE hybrids produced by SE substitution(s) using the N-terminal trunk and/or inhibitory loops of StCYS5, PpCYS or CsCYS (Table 3, this study), or *E. coli*-produced cystatins representative of the plant cystatin protein family (Table 2, this study). Data are expressed as *K*_i (papain)_ values for wild-type SlCYS8 relative to *K*_i (papain)_ values for the different cystatins or cystatin hybrids. *K*_i_ ratios were inferred from Table I of Goulet al. 2008 (single mutants); Figure 4, this study (SE hybrids), and Table 2, this study (original cystatins). The vertical red line highlights the reference *K*_i (papain)_ ratio of 1 as calculated for wild-type SlCYS8, and the blue lines average ratios for the three cystatin variant datasets. SE, structural element; CV, coefficient of variation.

## DISCUSSION

Protein engineering approaches have been proposed by several groups to improve the inhibitory properties of plant cystatins against herbivorous pest digestive Cys proteases (van Wyk et al. 2016, Tremblay et al. 2019). These strategies typically involve point mutations at functionally relevant amino acid sites or phage display procedures to select improved variants produced by random mutagenesis in the inhibitory loops. Here, we explored the potential of structural element substitutions as an alternative to these approaches, using tomato SlCYS8 and the digestive Cys cathepsins of *L. decemlineata* as a protease–inhibitor model system. Our data confirm the usefulness of natural cystatins among plant taxa as a pool of discrete function-related structural elements for the design of stable and active cystatin variants. They also confirm the potential of structural element substitutions to improve the inhibitory efficiency of tomato SlCYS8 against Cys proteases, in line with the reported robustness of plant cystatin structures and the usefulness of these proteins as translational fusion partners or scaffolds for different applications of biotechnological value (Sainsbury et al. 2013, 2016, Kyle 2018).

Our underlying assumption for this work was that substituting the N-terminal trunk and/or inhibitory loops of a cystatin would represent a useful complement to current protein engineering approaches by allowing conformational changes on a length scale beyond that accessible to single mutations (Clark et al. 2009). Supporting this, SE substitutions as here implemented had a much greater impact on SlCYS8 inhibitory activities than our previously described approach involving single substitutions at positively selected amino acid sites (**Figure 6**). Much interestingly, SE hybrids showed average *K*_i (papain)_ values improved by 150% overall relative to wild-type SlCYS8, five times higher than the overall improvement rate observed with the site-directed mutagenesis approach. Likewise, potent single mutants such as P2L, P2M and P2F produced earlier exhibited inhibitory activity rates increased by–i.e. *IC*_50_ values decreased by– two- to threefold against *L. decemlineata* cathepsin L-like enzymes compared to wild-type SlCYS8 (Goulet et al. 2008). By comparison, most SE variants bearing one or two structural elements of StCYS5 or PpCYS here showed five- to tenfold inhibitory rate increases against these enzymes in non-saturating conditions (**Figure 5**). Overall, these observations point to the potential of SE substitutions as an effective way to derive functionally diverse cystatin variants from a plant cystatin template, and hence the potential of this approach as a valuable complement to current protein engineering strategies for cystatin molecular improvement.

An unsolved question at this point is the actual relevance of our new approach in practice considering the functional variability already observed among plant cystatin family members and the high inhibitory efficiency here measured for some of them. For instance, the inhibitory effect of potent SE hybrids like P-N1 or P-1 against papain was much stronger than the anti-papain activity of wild-type SlCYS8 but comparable to the inhibitory activity of PpCYS used as a donor cystatin for the two variants (**Figure 4**, **Table 2**). Similarly, StCYS5 and PpCYS both showed inhibitory values 10 times higher than SlCYS8 against *L. decemlineata* cathepsin L enzymes, comparable to the inhibitory values observed for the two most potent SE hybrids, S-12 and P-12, bearing the inhibitory loops of these donor cystatins (**Figure 2**, **Figure 5**). On the other hand, hybrid S-N1, with the N-terminal trunk and first inhibitory loop of StCYS5, exhibited very strong inhibitory activity against *L. decemlineata* cathepsin B-like enzymes, more than 15 times higher than, and approximately three times higher than, the inhibitory activities of SlCYS8 and StCYS5, respectively (**Figure 5**). These data, while leaving open the question of a comparative plus-value for the SE substitutions strategy to produce potent inhibitors of cathepsin L-like enzymes, suggest the potential of this approach to generate broad-spectrum cystatins that also inhibit alternative proteases naturally recalcitrant to cystatin inhibition (Zhu-Salzman and Zeng 2015), such as those upregulated in *L. decemlineata* to sustain leaf consumption and larval growth (Rasoolizadeh et al. 2016b).

Additional studies will now be welcome to compare the plant protective effects of potent SE variants like hybrids S-N1, S-12 and P-12 with the protective effects of potent natural cystatins like StCYS5 and PpCYS or those of SlCYS8 single variant ^P2V^SlCYS8, an improved but still moderately efficient inhibitor (*see* **Figure 6**) shown to negatively alter leaf consumption and growth of *L. decemlineata* larvae (Rasoolizadeh et al. 2016b). Studies will also be welcome to assess the inhibitory potential of SE hybrids integrating N-terminal trunk and inhibitory loop(s) of different cystatin donors, given the specific contributions of these structural elements to the Cys protease–cystatin complex. Studies will be welcome, finally, to measure the impact of a fourth ‘structural element’, the central fold supporting the N-terminal trunk and two inhibitory loops, on the inhibitory activity of plant cystatins. The strong inhibitory potency of hybrid S-N1 against *L. decemlineata* cathepsin B-like enzymes compared to wild-type StCYS5 used as a donor, or the occurrence of positively selected amino acids presumably influencing protease inhibition in the α-helix and inter-loop region of plant cystatins (Kiggundu et al. 2006), point to a possible impact of this structural element on the inhibitory efficiency of the protein. The central fold of cystatins is not directly involved in protease inhibition but still might represent a valuable target for plant cystatin engineering given its possible effects on the spatial orientation and stability of the three functional elements.

## EXPERIMENTAL PROCEDURES

### Cys protease–cystatin docking simulations

Enzyme–inhibitor docking simulations were performed for tomato cystatins SlCYS8 (GenBank Accession No. AF198390) and SlCYS9 (GenBank NP001275067), soybean cystatin GmCYS2 (GenBank AAA97906), rice cystatin OsCYS1 (GenBank NP001044550) and corn cystatin ZmCYS1 (GenBank NP001105295) interacting with papain, human cathepsin L and *L. decemlineata* IntD4 (GenBank EF154436) considered as target Cys proteases. Simulations were performed using the Z-Dock algorithm of Discovery Studio (Accelrys Software Inc.) after inferring structure homology models for IntD4 and the five plant cystatins. Twenty tentative models were built using Modeller, v. 9.7 (Sali and Blundell 1993, Marti-Renom et al. 2000), with the crystal structure of human cathepsin L (PDB 1SC8) as a template for the insect protease (Sainsbury et al. 2012b) and the NMR structure of oryzacystatin I (PDB 1EQK) as a template for the cystatins. Stereochemical quality of the models was compared to their template structures with the Procheck program, v.3.5.4 (http://www.ebi.ac.uk/thornton-srv/software/PROCHECK/) (Laskowski et al. 1993) and the best models were selected for further analyses. Docking simulations with models of the five cystatins were produced for papain (PDB 9PAP), cathepsin L (1SC8) and IntD4 (Vorster et al. 2015) using the Z-Dock algorithm to generate 2,000 tentative poses for the resulting complexes. Top-ranking poses, based on the Z-score (Chen et al. 2003), were compared with the solved crystal structure of human stefin B in complex with papain (PDB 1STF) to confirm the relative binding positions and orientations of the proteins in the predicted complexes. Five tentative complexes were chosen for each protease–cystatin combination and refined through energy minimization using the R-Dock algorithm (Li et al. 2003). Interacting residues and interaction (binding) energies were inferred for the top-ranking models. Normal distribution tests were performed on calculated data using the Shapiro-Wilk test of normality (Shapiro and Wilk 1965), followed by a F-test to compare the variances of two samples structural element combinations (N-terminal trunk vs the first inhibitory loop; N-terminal trunk vs the second inhibitory loop; and first loop vs the second loop) from normal populations. An alpha threshold of 5% was used for statistical significance.

### Representative plant cystatins

Phylogenetic inferences were performed with the MEGA6 software, v.6.06 (Tamura et al. 2013), using plant cystatin sequences available in the NCBI protein database (http://ncbi.nlm.nih.gov/). Non-redundant cystatin sequences were retrieved from the Viridiplantae domain of the database using the in-built Protein Blast tool, with the sequence of tomato SlCYS8 as a query sequence (GenBank Accession No. AF198390.1; gi|6671196). Amino acid sequences including at least 90% of a full cystatin, here corresponding to 262 non-redundant NCBI accessions, were used as a starting point. A multiple sequence alignment was generated using the MUSCLE algorithm (Edgar 2004), from which a maximum likelihood tree was calculated based on the “JTT” amino acid substitution model (Jones et al. 1992) (**Supplementary Figure 2**). Fifty-seven ‘representative’ cystatins were identified by restricting the selection (i) to one sequence among highly similar sequences (>95% identity), and (ii) to one cystatin per branch of the phylogenetic tree. A subset of 30 cystatins was taken from this sample for the functional analyses, in such a way as to maximize sequence variability among the selected cystatins at amino acid positions expected to physically interact with the target proteases (Vorster et al. 2015).

### Recombinant cystatins

All cystatins were produced in *E. coli*, strain BL21 as described previously (Goulet et al. 2008), using the GST gene fusion system for heterologous expression and affinity purification (GE Healthcare). DNA templates for the cystatins were synthesized as g-blocks (IDT) including GoldenGate BSAI cloning sites on both sides of the cystatin coding region (Sainsbury et al. 2012b). DNA coding sequences for the original cystatins corresponded to those sequences reported in GenBank (as listed in **Table 2**). DNA sequences for the SE hybrids were designed as described in the Results (**Figure 3**), with structural elements from donor cystatins potato StCYS5, *P. patens* PpCYS or cucumber CsCYS replacing the N-terminal trunk (amino acids 1– 13, SlCYS8-numbering), first inhibitory loop (amino acids 44–55) and/or second inhibitory loop (amino acid 70–84) of tomato SlCYS8 (**Supplementary Table 2**). G-blocks were inserted in a modified version of the pGEX-3X expression vector (GE Healthcare) using the Golden Gate DNA shuffling method of Engler et al. (2009), downstream of a ‘GST–factor X_a_ cleavage site’ coding sequence (Sainsbury et al. 2012b). The GST tag was removed by cleavage with bovine factor X_a_, according to the manufacturer’s specifications (Novagen). Cystatin products showing fragmentation, as assessed by 15% (w/v) SDS-PAGE, were not considered further for the functional analyses. The purified cystatins were quantified by densitometric analysis of Coomassie blue-stained polyacrylamide slab gels following 12% (w/v) SDS-PAGE, using three technical replicates and bovine serum albumin (Sigma-Aldrich) as a protein standard.

### Test proteases

Papain (E.C.3.4.22.2, from papaya latex) was purchased from Sigma-Aldrich. Colorado potato beetle (*L. decemlineata*) proteases were extracted from the midgut of fourth instars reared on greenhouse-grown potato plants, cv. Norland, as described previously (Goulet et al. 2008). Two-spotted spider mite (*T. urticae*) proteases were obtained from a laboratory colony reared in greenhouse on common bean (*Phaseolus vulgaris*). Whole mites were ground in liquid nitrogen, the resulting powder kept on ice for 10 min after resuspension in 50 mM Tris-HCl buffer, pH 7.0, and the whole mixture centrifuged at 4°C for 10 min at 20 000 *g*. The pellet was discarded, and the supernatant used as a source of digestive proteases for the protease inhibitory assays. Papain and soluble proteins in the arthropod crude extracts were assayed according to Bradford (1976), with bovine serum albumin as a protein standard.

### *K*_i (papain)_ value determinations

*K*_i (papain)_ values for the cystatin variants were determined by the monitoring of substrate hydrolysis progress curves (Salvesen and Nagase 1989), based on the linear equation of Henderson (1972). Papain activity was monitored in 50 mM Tris-HCl, pH 7.0, using the synthetic peptide substrate *Z*-Phe–Arg-methylcoumarin (MCA) (Sigma-Aldrich). Hydrolysis was allowed to proceed at 25°C in reduced conditions (10 mM L-cysteine) with the substrate in large excess, after adding (or not) recombinant cystatins dissolved in a minimal volume of reaction buffer. Papain activity was monitored using a Synergy H1 fluorimeter (BioTek), using an excitation filter of 360 nm and an emission filter of 450 nm. *K*_i_ values were calculated using the experimentally determined *K*_i(app)_ and *K*_m_ values, based on the following equation: *K*_i_ = *K*_i(app)_ / (1 + [S] / *K*_m_). A *K*_m_ value of 93.6 μM was determined for papain under our assay conditions.

### Arthropod protease assays

Cys cathepsin activities in the arthropod protein extracts were assayed in 0.2 M NaH_2_PO_4_/0.2 M Na_2_HPO_4_ phosphate buffer, pH 6.5, using the synthetic peptide substrates *Z*-Phe–Arg-MCA for cathepsin L-like activities and *Z*-Arg–Arg-MCA for cathepsin B-like activities. Hydrolysis was allowed to proceed in reduced conditions (10 mM L-cysteine) for 10 min at 25°C, with ~5-6 ng of arthropod protein per μl in the reaction mixture and the peptide substrate added in large excess. Cystatins dissolved in a minimal volume of reaction buffer were added to the reaction mixture for the inhibitory assays. Proteolytic activity was monitored using a Synergy H1 fluorimeter (BioTek), with excitation and emission filters of 360 nm and 450 nm, respectively.

## ACKNOWLEDGEMENTS

This work was supported by a Discovery grant from the Natural Science and Engineering Research Council of Canada.

## SUPPLEMENTARY MATERIAL

**Supplementary Figure 1** Complement to Table 1 : Amino acid sequence alignments of plant cystatins SlCYS8, SlCYS9, GmCYS2, OsCYS1 and ZmCYS1 highlighting residues predicted to interact with papain, human cathepsin L or *L. decemlineata* IntD4.

**Supplementary Figure 2** Complement to Table 2 : Maximum likelihood phylogenetic tree generated for 262 plant cystatin amino acid sequences available in the NCBI protein database.

**Supplementary Figure 3** Inhibition of *T. urticae Z*-Phe–Arg-MCA-hydrolyzing (cathepsin L-like) enzymes by the SlCYS8 SE hybrids.

**Supplementary Table 1** Complement to Table 1: Interaction binding energies inferred *in silico* for model Cys proteases papain, human cathepsin L and *L. decemlineata* IntD4 interacting with N-terminal trunk, Loop 1 and Loop 2 amino acids of tomato cystatins SlCYS8 and SlCYS9, rice cystatin OsCYS1, soybean cystatin GmCYS2 and corn cystatin ZmCYS1

**Supplementary Table 2** Primary sequences of recipient cystatin SlCYS8, donor cystatins StCYS5, PpCYS and CsCYS, and SlCYS8 SE hybrids bearing one, two or three structural elements of StCYS5, PpCYS or CsCYS.

## LITERATURE CITED

Arai S, Matsumoto I, Emori Y, Abe K (2002) Plant seed cystatins and their target enzymes of endogenous and exogenous origin. J Agric Food Chem 50, 6612–6617.

Benchabane M, Schlüter U, Vorster J, Goulet M-C, Michaud D (2010) Plant cystatins. Biochimie 92, 1657–1666.

Bradford MM (1976) A rapid and sensitive method for the quantitation of microgram quantities of protein utilizing the principle of protein-dye binding. Anal Biochem 72, 248–254.

Broadway RM (2001) The adaptation of insects to protease inhibitors, in: Michaud, D. (Ed.), Recombinant protease inhibitors in plants. CRC Press, Boca Raton FL, pp. 80–88.

Chen PJ, Senthilkumar R, Jane WN, He Y, Tian Z, Yeh KW (2014) Transplastomic *Nicotiana benthamiana* plants expressing multiple defence genes encoding protease inhibitors and chitinase display broad-spectrum resistance against insects, pathogens and abiotic stresses. Plant Biotechnol J 12, 503–515.

Chen R, Li L, Weng Z (2003) ZDOCK: an initial-stage protein-docking algorithm. Proteins 52, 80–87.

Cingel A, Savić J, Vinterhalter B, Vinterhalter D, Kostić M, Jovanović DS, Smigocki A, Ninković S (2015) Growth and development of Colorado potato beetle larvae, *Leptinotarsa decemlineata*, on potato plants expressing the oryzacystatin II proteinase inhibitor. Transgenic Res 24, 729–740.

Clark LA, Boriack-Sjodin PA, Day E, Eldredge J, Fitch C, Jarpe M, Miller S, Li Y, Simon K, van Vlijmen WT (2009) Prot Eng Des Select 22, 93–101.

Cloutier C, Jean C, Fournier M, Yelle S, Michaud D (2000) Adult Colorado potato beetles, *Leptinotarsa decemlineata* compensate for nutritional stress on oryzacystatin I transgenic potato plants by hypertrophic behavior and over-production of insensitive proteases. Arch Insect Physiol Biochem 44, 69–81.

Cloutier C, Fournier M, Jean C, Yelle S, Michaud D (1999) Growth compensation and faster development of Colorado potato beetle (Coleoptera: Chrysomelidae) feeding on potato foliage expressing oryzacystatin I. Arch Insect Biochem Physiol 40, 69–79.

Edgar RC (2004) MUSCLE: Multiple sequence alignment with high accuracy and high-throughput. Nucl Acids Res 32, 1792–1797.

Engler C, Gruetzner R, Kandzia R, Marillonnet S (2009) Golden gate shuffling: a one-pot DNA shuffling method based on type IIs restriction enzymes. PLoS One. 4, e5553.

Goulet M-C, Dallaire C, Vaillancourt L-P, Khalf M, Badri AM, Preradov A, Duceppe M-O, Goulet C, Cloutier C, Michaud D (2008) Tailoring the specificity of a plant cystatin toward herbivorous insect digestive cysteine proteases by single mutations at positively selected amino acid sites. Plant Physiol 146, 1010–1019.

Grosse-Holz F, Madeira L, Zahid MA, Songer M, Kourelis J, Fesenko M, Ninck S, Kaschani F, Kaiser M, van der Hoorn RAL (2018) Three unrelated protease inhibitors enhance accumulation of pharmaceutical recombinant proteins in *Nicotiana benthamiana*. Plant Biotechnol J 16, 1797–1810

Henderson PJF (1972) A linear equation that describes the steady-state kinetics of enzymes and subcellular particles interacting with tightly bound inhibitors. Biochem J 127, 321–333.

Hwang JE, Hong JK, Je JH, Lee KO, Kim DY, Lee SY, Lim CO (2009) Regulation of seed germination and seedling growth by an *Arabidopsis* phytocystatin isoform, *AtCYS6*. Plant Cell Rep 28, 1623–1632.

Jones DT, Taylor WR, Thornton JM (1992) The rapid generation of mutation data matrices from protein sequences. Comput Appl Biosci 8, 275–282.

Jutras PV, Grosse-Holz F, Kaschani F, Kaiser M, Michaud D, van der Hoorn RAL (2019) Activity based proteomics reveals nine target proteases for the recombinant protein-stabilizing inhibitor *Sl*CYS8 in *Nicotiana benthamiana*. Plant Biotechnol J 17, 1670–1678.

Jutras PV, Goulet M-C, Lavoie P-O, D’Aoust M-A, Sainsbury F, Michaud D (2018) Recombinant protein susceptibility to proteolysis in the plant cell secretory pathway is pH-dependent. Plant Biotechnol J 16, 1928–1938.

Jutras PV, Marusic C, Lonoce C, Deflers C, Goulet M-C, Benvenuto E, Michaud D, Donini M (2016) An accessory protease inhibitor to increase the yield and quality of a tumour-targeting mAb in *Nicotiana benthamiana* leaves. PLoS One 11, e0167086.

Kiggundu A, Goulet M-C, Goulet C, Dubuc JF, Rivard D, Benchabane M, Pépin G, van der Vyver C, Kunert K, Michaud D (2006) Modulating the proteinase inhibitory profile of a plant cystatin by single mutations at positively selected amino acid sites. Plant J 48, 403–413.

Kyle S (2018) Affimer proteins : Theranostics of the future? Trends Biochem Sci 43, 230–232.

Laskowski RA, MacArthur MW, Moss DS, Thornton JM (1993) *PROCHECK*: a program to check the stereochemical quality of protein structures. J Appl Cryst 26, 283–291.

Lecardonnel A, Chauvin L, Jouanin L, Beaujean A, Prévost G, Sangwan-Norreel B (1999) Effects of rice cystatin I expression in transgenic potato on Colorado potato beetle larvae. Plant Sci 140, 71–79.

Li L, Chen R, Weng Z (2003) RDOCK: refinement of rigid-body protein docking predictions. Proteins 53, 693–707.

Lima AM, dos Reis SP, de Souza CRB (2015) Phytocystatins and their potential to control plant diseases caused by fungi. Prot Pept Lett 22, 104–111.

Ljunggren A, Redzynia I, Alvarez-Fernandez M, Abrahamson M, Mort JS, Krupa JC, Jaskolski M, Bujacz G (2007) Crystal structure of the parasite protease inhibitor chagasin in complex with a host target cysteine protease. J Mol Biol 371, 137–153.

Macedo MLR, de Oliveira CFR, Costa PM, Castelhano EC, Silva-Filho MC (2015) Adaptive mechanisms of insect pests against plant protease inhibitors and future prospects related to crop protection: A review. Prot Pept Lett 22, 149–163.

Marcos E, Basanta B, Chidyausiku TM, Tang Y, Oberdorfer G, Liu G, Swapna GVT, Guan R, Silva DA, Dou J, Pereira JH, Xiao R, Sankaran B, Zwart PH, Montelione GT, Baker D (2017) Principles for designing proteins with cavities formed by curved ß sheets. Science 355, 201–206.

Marti-Renom MA, Stuart A, Fiser A, Sánchez R, Melo F, Sali A (2000) Comparative protein structure modeling of genes and genomes. Annu Rev Biophys Biomol Struct 29, 291–325.

Martinez M, Santamaria ME, Díaz-Mendoza M, Arnaiz A, Carrillo L, Ortego F, Díaz I (2016) Phytocystatins: Defense proteins against phytophagous insects and acari. Int J Mol Sci 17, 1747.

Munger A, Simon M-A, Khalf M, Goulet M-C, Michaud D (2015) Cereal cystatins delay sprouting and nutrient loss in tubers of potato, *Solanum tuberosum*. BMC Plant Biol 15, 296.

Munger A, Coenen K, Cantin L, Goulet C, Vaillancourt L-P, Goulet M-C, Tweddell R, Sainsbury F, Michaud D (2012) Beneficial ‘unintended effects’ of a cereal cystatin in transgenic lines of potato, *Solanum tuberosum*. BMC Plant Biol 12, 198.

Nagata K, Kudo N, Abe K, Arai S, Tanokura M (2000) Three-dimensional solution structure of oryzacystatin-I, a cysteine proteinase inhibitor of the rice, *Oryza sativa* L. japonica. Biochemistry 39, 14753–14760.

Quain MD, Makgopa ME, Márquez-García B, Comadira G, Fernendez-Garcia N, Olmos E, Schnaubelt D, Kunert KJ, Foyer CH (2014) Ectopic phytocystatin expression leads to enhanced drought stress tolerance in soybean (*Glycine max*) and *Arabidopsis thaliana* through effects on strigolactone pathways and can also result in improved seed traits. Plant Biotechnol J 12, 903–913.

Rasoolizadeh A, Goulet M-C, Sainsbury F, Cloutier C, Michaud D (2016a) Single substitutions to closely related amino acids contribute to the functional diversification of an insect-inducible, positively selected plant cystatin. FEBS J 283, 1323–1335.

Rasoolizadeh A, Munger A, Goulet M-C, Sainsbury F, Cloutier C, Michaud D (2016b) Functional proteomics-aided selection of protease inhibitors for herbivore insect control. Sci Rep 6, 38827.

Sainsbury F, Jutras PV, Vorster J, Goulet M-C, Michaud D (2016) A chimeric affinity tag for efficient expression and chromatographic purification of heterologous proteins from plants. Front Plant Sci 7, 141.

Sainsbury F, Varennes-Jutras P, Goulet M-C, D’Aoust M-A, Michaud D (2013) Tomato cystatin *Sl*CYS8 as a stabilizing fusion partner for human serpin expression in plants. Plant Biotechnol J 11, 1058–1068.

Sainsbury F, Benchabane M, Goulet M-C, Michaud D (2012a) Multimodal protein constructs for herbivore insect control. Toxins 4, 455–475.

Sainsbury F, Rhéaume A-J, Goulet M-C, Vorster J, Michaud D (2012b) Discrimination of differentially inhibited cysteine proteases by activity-based profiling using cystatin variants with tailored specificities. J Proteome Res 11, 5983–5993.

Sali A, Blundell TL (1993) Comparative protein modelling by satisfaction of spatial restraints. J Mol Biol 234, 779–815.

Salvesen G, Nagase H (1989) Inhibition of proteolytic enzymes. In Proteolytic enzymes. A practical approach (Bond J.S. and Beynon, R., eds.). New York: IRL Press, pp. 83–104.

Schlüter U, Benchabane M, Munger A, Kiggundu A, Vorster J, Goulet M-C, Cloutier C, Michaud D (2010) Recombinant protease inhibitors for herbivore pest control: a multitrophic perspective. J Exp Bot 61, 4169–4183.

Shapiro SS, Wilk MB (1965) An analysis of variance test for normality (complete samples). Biometrika 52, 591611.

Shukla P, Subhashini M, Singh NV, Ahmed I, Trishla S, Kirti PB (2016) Targeted expression of cystatin restores fertility in cysteine protease induced male sterile tobacco plants. Plant Sci 246, 52–61.

Tamura K, Stecher G, Peterson D, Filipski A, Kumar S (2013) MEGA6 : Molecular evolutionary genetics analysis version 6.0. Mol Biol Evol 30, 2725–2729.

Tan Y, Li M, Yang Y, Sun X, Wang N, Liang B, Ma F (2017a) Overexpression of *MpCYS4*, a phytocystatin gene from *Malus prunifolia* (Willd.) Borkh., enhances stomatal closure to confer drought tolerance in transgenic *Arabidopsis* and apple. Front Plant Sci 8, 33.

Tan Y, Yang Y, Li C, Liang B, Li M, Ma F (2017b) Overexpression of *MpCYS4,* a phytocystatin gene from *Malus prunifolia* (Willd.) Borkh., delays natural and stress-induced leaf senescence in apple. Plant Physiol Biochem 115, 219–228.

Tan Y, Wei X, Wang P, Sun X, Li M, Ma F (2016) A phytocystatin gene from *Malus prunifolia* (Willd.) Borkh., *MpCYS5*, confers salt tolerance and functions in endoplasmic reticulum stress response in *Arabidopsis*. Plant Mol Biol Rep 34, 62–75.

Tiede C, Tang AAS, Deacon SE, Mandal U, Nettleship JF, Owen RL, George SE, Harrison DJ, Owens RJ, Tomlinson DC, McPherson MJ (2014) Adhiron: a stable and versatile peptide display scaffold for molecular recognition applications. Protein Eng Des Select 27, 145–155.

Tremblay J, Goulet M-C, Michaud D (2019) Recombinant cystatins in plants. Biochimie 166, 184–193.

Turk V, Bode W (1991) The cystatins: protein inhibitors of cysteine proteinases. FEBS Lett 285, 213–219.

Urwin PE, Atkinson HJ, Waller DA, McPherson MJ (1995) Engineered oryzacystatin-I expressed in transgenic hairy roots confers resistance to *Globodera pallida*. Plant J 8, 121–131.

van Wyk SG, Kunert KJ, Cullis CA, Piullay P, Makgopa ME, Schlüter U, Vorster BJ (2016) The future of cystatin engineering. Plant Sci 246, 119–127.

Vorster BJ, Tastan Bishop Ö, Schlüter U, Coetzer N, Michaud D (2010) New insights towards the understanding of phytocystatin–papain interactions. Aspects Appl Biol 96, 403–408.

Vorster J, Rasoolizadeh A, Goulet M-C, Cloutier C, Sainsbury F, Michaud D (2015) Positive selection of digestive Cys proteases in herbivorous Coleoptera. Insect Biochem Mol Biol 65, 10–19.

Zhu-Salzman K, Zeng R (2015) Insect response to plant defensive protease inhibitors. Annu Rev Entomol 60, 233–252.

