## Supplementary Material for "Harnessing the functional diversity of plant cystatins to design inhibitor variants highly active against herbivorous arthropod digestive proteases"

**Supplementary Figures 1–3 | Supplementary Tables 1 and 2**

Residues interacting with papain

```
                10         20         30         40         50
            ....|....| ....|....| ....|....| ....|....| ....|....|
SlCYS8      .SPNPGGITN VPFP..NLPQ FKDLARFAVQ DYNKKENAHL EFVENLNVKE
SlCYS9      .MATLGGVHD SHGSSQNSDE IHSLAKFAVD EHNKKENAMI ELARVVKAQE
GmCYS2      .VQELGGITD VHGAA.NSVE INNLARFAVE EQNKRENSVL EFVRVISAKQ
OsCYS1      GGPVLGGVEP V.GNE.NDLH LVDLARFAVT EHNKKANSLL EFEKLVSVKQ
ZmCYS1      AGMLAGGIKD VPANE.NDLQ LQELARFAVN EHNQKANALL GFEKLVKAKT

                60         70         80         90        100
            ....|....| ....|....| ....|....| ....|....| ....|....|
SlCYS8      QVVAGIIYYI TLVATDAGKK KIYETKILVK GWENFKEVQE FKLVGDATKP
SlCYS9      QTVAGKLHHL TLEVMDAGKK KLYEAKVWVK PWLNFKELQE FKHVEDVPTF
GmCYS2      QVVAGVNYYI TLEAKDGLIK NEYEAKVWVR EWLNSKELLE FKPVNVSSTP
OsCYS1      QVVAGTLYYF TIEVKEGDAK KLYEAKVWEK PWMDFKELQE FKPVDASANA
ZmCYS1      QVVAGTMYYL TIEVKDGEVN KLYEAKVWEK PWENFKQLQE FKPVEEGASA
```

Residues interacting with human cathepsin L

```
                10         20         30         40         50
            ....|....| ....|....| ....|....| ....|....| ....|....|
SlCYS8      .SPNPGGITN VPFP..NLPQ FKDLARFAVQ DYNKKENAHL EFVENLNVKE
SlCYS9      .MATLGGVHD SHGSSQNSDE IHSLAKFAVD EHNKKENAMI ELARVVKAQE
GmCYS2      .VQELGGITD VHGAA.NSVE INNLARFAVE EQNKRENSVL EFVRVISAKQ
OsCYS1      GGPVLGGVEP V.GNE.NDLH LVDLARFAVT EHNKKANSLL EFEKLVSVKQ
ZmCYS1      AGMLAGGIKD VPANE.NDLQ LQELARFAVN EHNQKANALL GFEKLVKAKT

                60         70         80         90        100
            ....|....| ....|....| ....|....| ....|....| ....|....|
SlCYS8      QVVAGIIYYI TLVATDAGKK KIYETKILVK GWENFKEVQE FKLVGDATKP
SlCYS9      QTVAGKLHHL TLEVMDAGKK KLYEAKVWVK PWLNFKELQE FKHVEDVPTF
GmCYS2      QVVAGVNYYI TLEAKDGLIK NEYEAKVWVR EWLNSKELLE FKPVNVSSTP
OsCYS1      QVVAGTLYYF TIEVKEGDAK KLYEAKVWEK PWMDFKELQE FKPVDASANA
ZmCYS1      QVVAGTMYYL TIEVKDGEVN KLYEAKVWEK PWENFKQLQE FKPVEEGASA
```

Residues interacting with *L. decemlineata* Intestain D4

```
                10         20         30         40         50
            ....|....| ....|....| ....|....| ....|....| ....|....|
SlCYS8      .SPNPGGITN VPFP..NLPQ FKDLARFAVQ DYNKKENAHL EFVENLNVKE
SlCYS9      .MATLGGVHD SHGSSQNSDE IHSLAKFAVD EHNKKENAMI ELARVVKAQE
GmCYS2      .VQELGGITD VHGAA.NSVE INNLARFAVE EQNKRENSVL EFVRVISAKQ
OsCYS1      GGPVLGGVEP V.GNE.NDLH LVDLARFAVT EHNKKANSLL EFEKLVSVKQ
ZmCYS1      AGMLAGGIKD VPANE.NDLQ LQELARFAVN EHNQKANALL GFEKLVKAKT

                60         70         80         90        100
            ....|....| ....|....| ....|....| ....|....| ....|....|
SlCYS8      QVVAGIIYYI TLVATDAGKK KIYETKILVK GWENFKEVQE FKLVGDATKP
SlCYS9      QTVAGKLHHL TLEVMDAGKK KLYEAKVWVK PWLNFKELQE FKHVEDVPTF
GmCYS2      QVVAGVNYYI TLEAKDGLIK NEYEAKVWVR EWLNSKELLE FKPVNVSSTP
OsCYS1      QVVAGTLYYF TIEVKEGDAK KLYEAKVWEK PWMDFKELQE FKPVDASANA
ZmCYS1      QVVAGTMYYL TIEVKDGEVN KLYEAKVWEK PWENFKQLQE FKPVEEGASA
```

**Supplementary Figure 1** Complement to Table 1 : Amino acid sequence alignments of cystatins SlCYS8, SlCYS9, GmCYS2, OsCYS1 and ZmCYS1 highlighting residues predicted to interact with papain (in blue), human cathepsin L (in green) or *L. decemlineata* IntD4 (in yellow).

---- See attached EPS file for figure display ----

**Supplementary Figure 2** Complement to Table 2 : Maximum likelihood phylogenetic tree generated for 262 plant cystatin amino acid sequences available in the NCBI protein database. The tree was calculated based on the 'JTT' amino acid substitution model of Jones et al. (1992) after generating a multiple sequence alignment using the MUSCLE algorithm (Edgar 2004). Red boxes highlight the 20 cystatins used for protease inhibitory assays.

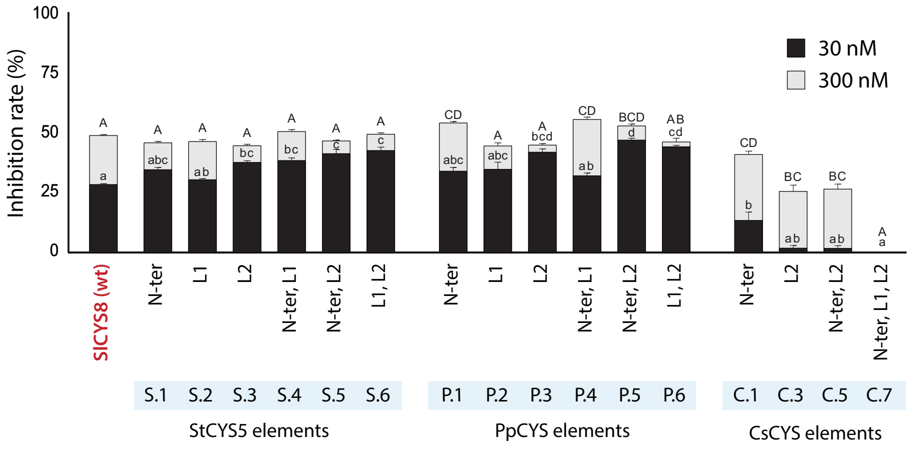


**Supplementary Figure 3** Inhibition of *T. urticae* Z-Phe–Arg-MCA-hydrolyzing (cathepsin L-like) enzymes by the SlCYS8 SE hybrids. Data are expressed as relative inhibitory rates compared to the inhibitory rate measured E-64 (100%). Inhibitory assays were conducted with limiting (30 nM) or excess (300 nM) concentrations of cystatin. Each bar is the mean of three independent (biological) replicates ± SE. Different letters (lower case for the limiting concentrations, capital for the excess concentrations) indicate significantly different inhibitory rates among cystatin hybrids (post-ANOVA Tuckey's mean comparison test, with an alpha value of 5%). S, P and C stand for StCYS5, PpCYS and CsCYS, respectively; N, 1 and 2 for the N-terminal trunk, first inhibitory loop and second inhibitory loop of the donor cystatin.

**Supplementary Table 1** Complement to Table 1: Interaction binding energies inferred *in silico* for model Cys proteases papain, human cathepsin L and *L. decemlineata* IntD4 interacting with N-terminal trunk, Loop 1 and Loop 2 amino acids of tomato cystatins SlCYS8 and SlCYS9, rice cystatin OsCYS1, soybean cystatin GmCYS2 and corn cystatin ZmCYS1.

**SlCYS8**

| | | Interaction energy (kcal/mol) | | |
|---|---|---|---|---|
| | Residue | Papain | Cathepsin L | Intestain D4 |
| N-ter | SER 1 | -21,90 | -204,86 | -209,55 |
| | PRO 2 | -22,18 | -18,38 | -2,27 |
| | ASN 3 | -35,08 | -50,72 | -45,55 |
| | PRO 4 | -28,44 | -40,11 | -28,34 |
| | GLY 5 | -20,82 | -23,66 | -30,81 |
| | GLY 6 | | -8,37 | -14,64 |
| | ILE 7 | -14,71 | | |
| | THR 8 | | -29,86 | |
| | VAL 10 | | -10,48 | |
| | GLN 17 | | | -2,08 |
| | PHE 18 | 2,10 | | |
| | | -141,02 | -386,44 | -333,26 |
| Loop 1 | GLU 47 | | 120,83 | |
| | GLN 48 | | -35,93 | |
| | VAL 49 | | -30,95 | |
| | VAL 50 | -11,68 | -48,30 | -15,45 |
| | ALA 51 | -22,75 | -30,73 | -28,76 |
| | GLY 52 | -23,96 | -38,27 | -25,14 |
| | ILE 53 | -23,11 | -33,80 | -10,28 |
| | ILE 54 | -17,17 | -7,47 | -4,77 |
| | TYR 55 | -7,24 | -12,98 | |
| | TYR 56 | | -15,85 | |
| | | -105,90 | -133,45 | -84,40 |
| Loop 2 | LYS 73 | -8,19 | -104,79 | |
| | LEU 75 | -25,20 | -17,46 | -18,21 |
| | VAL76 | -24,88 | | |
| | LYS 77 | -53,42 | -66,49 | -112,76 |
| | GLY 78 | -20,91 | -30,65 | -4,98 |
| | TRP 79 | -43,81 | -46,39 | -12,75 |
| | GLU 80 | -56,66 | -45,91 | 27,96 |
| | ASN 81 | -41,38 | -41,37 | -20,98 |
| | PHE 82 | -20,80 | -9,10 | |
| | LYS 83 | -16,27 | | |
| | GLU 84 | -49,60 | -28,78 | -98,38 |
| | GLN 86 | -24,48 | -17,66 | |
| | | -385,60 | -408,61 | -240,10 |

**SlCYS9**

| | | Interaction energy (kcal/mol) | | |
|---|---|---|---|---|
| | Residue | Papain | Cathepsin L | Intestain D4 |
| N-ter | MET 1 | -28,12 | -206,91 | -182,80 |
| | ALA 2 | -12,42 | -28,85 | -27,04 |
| | THR 3 | -34,68 | -82,95 | -30,27 |
| | LEU 4 | -26,57 | -18,30 | -28,08 |
| | GLY 5 | -21,15 | -17,27 | -11,68 |
| | GLY 6 | | -14,15 | -15,17 |
| | VAL 7 | -10,70 | -20,54 | |
| | HIS 8 | | | |
| | GLU 19 | | | -7,15 |
| | | -133,65 | -388,98 | -302,18 |
| Loop 1 | GLU 49 | -81,49 | 76,24 | |
| | GLN 50 | | -22,78 | |
| | THR 51 | -16,72 | -38,82 | -23,42 |
| | VAL 52 | -18,30 | -33,16 | -10,76 |
| | ALA 53 | -16,69 | -20,40 | -17,98 |
| | GLY 54 | -10,64 | -41,46 | -16,78 |
| | LYS 55 | -10,57 | -224,96 | -18,61 |
| | LEU 56 | -25,66 | -9,73 | -4,60 |
| | HIS 58 | | -12,20 | |
| | | -180,08 | -327,27 | -92,15 |
| Loop 2 | TRP 77 | -34,28 | -19,88 | -18,93 |
| | VAL 78 | -13,86 | | |
| | LYS 79 | -112,50 | -77,85 | -124,41 |
| | PRO 80 | -12,73 | -25,72 | -11,54 |
| | TRP 81 | -46,16 | -59,20 | -27,69 |
| | LEU 82 | -39,47 | -24,58 | -28,35 |
| | ASN 83 | -32,86 | -28,83 | |
| | PHE 84 | -15,24 | -20,32 | |
| | LYS 85 | -12,49 | | |
| | GLU 86 | -63,05 | -3,34 | -60,77 |
| | GLN 88 | -18,96 | -14,80 | |
| | | -401,60 | -274,52 | -271,70 |

**Supplementary Table 1** Cont'd

| **OsCYS1** | | Interaction Energy (kcal/mol) | | |
|---|---|---|---|---|
| | Residue | Papain | Cathepsin L | Intestain D4 |
| N-ter | GLY 1 | -30,33 | -235,75 | -216,11 |
| | GLY 2 | -30,07 | -27,89 | -9,32 |
| | PRO 3 | -11,65 | -36,53 | -17,06 |
| | VAL 4 | -33,24 | -13,01 | -22,99 |
| | LEU 5 | -45,87 | -19,60 | -30,55 |
| | GLY 6 | | -51,68 | -28,90 |
| | GLY 7 | | -35,31 | -16,16 |
| | | -151,16 | -419,76 | -341,09 |
| L1 | GLN 48 | | -24,68 | |
| | GLN 49 | | -29,81 | -31,28 |
| | VAL 50 | -27,35 | -35,28 | |
| | VAL 51 | -17,10 | -44,36 | -11,14 |
| | ALA 52 | -28,99 | -39,44 | -15,48 |
| | GLY 53 | -21,76 | -24,66 | -19,91 |
| | THR 54 | -35,46 | -17,90 | -26,18 |
| | LEU 55 | -19,09 | -19,10 | |
| | TYR 56 | -20,21 | -19,78 | |
| | TYR 57 | | -18,25 | |
| | | -169,97 | -273,26 | -103,98 |
| L2 | LYS 74 | | -53,62 | |
| | VAL 75 | -7,61 | | |
| | TRP 76 | -31,21 | -29,39 | -21,25 |
| | GLU 77 | -32,47 | 21,42 | -20,32 |
| | LYS 78 | -83,83 | -103,06 | -102,75 |
| | PRO 79 | -23,96 | -25,67 | -7,82 |
| | TRP 80 | -30,66 | -36,49 | -21,16 |
| | MET 81 | -31,19 | -33,86 | -12,17 |
| | PHE 83 | -13,22 | -15,71 | -6,59 |
| | LYS 84 | -8,94 | | |
| | GLU 85 | -45,06 | 7,84 | -97,04 |
| | GLN 87 | -9,74 | -22,08 | |
| | | -317,89 | -290,62 | -289,11 |

| **GmCYS2** | | Interaction Energy (kcal/mol) | | |
|---|---|---|---|---|
| | Residue | Papain | Cathepsin L | Intestain D4 |
| N-ter | VAL 1 | -23,56 | -241,16 | -222,12 |
| | GLN 2 | -36,87 | -40,99 | -26,14 |
| | GLU 3 | -112,33 | -63,00 | -14,33 |
| | LEU 4 | -18,08 | -30,57 | -8,27 |
| | GLY 5 | -25,67 | -21,23 | -43,99 |
| | GLY 6 | | -3,99 | -11,91 |
| | ILE 7 | -21,29 | | -14,24 |
| | THR 8 | | -11,83 | |
| | GLU 18 | -95,98 | | -5,63 |
| | | -333,79 | -412,76 | -346,63 |
| L1 | GLN 48 | | -40,15 | |
| | GLN 49 | -35,75 | -23,03 | -26,91 |
| | VAL 50 | -18,15 | -37,44 | -5,83 |
| | VAL 51 | -15,64 | -29,73 | -18,19 |
| | ALA 52 | -27,80 | -30,86 | -42,93 |
| | GLY 53 | -19,20 | -41,75 | -24,88 |
| | VAL 54 | -19,41 | -33,31 | -24,32 |
| | ASN 55 | -35,73 | -29,56 | -22,72 |
| | TYR 56 | -17,02 | -16,78 | |
| | TYR 57 | | -20,97 | |
| | | -188,70 | -303,58 | -165,78 |
| L2 | VAL 75 | -5,71 | | |
| | TRP 76 | -28,87 | -33,99 | -16,45 |
| | VAL 77 | -27,05 | | |
| | ARG 78 | -97,08 | -123,87 | -12,34 |
| | GLU 79 | -141,77 | 51,79 | -14,30 |
| | TRP 80 | -34,58 | -45,96 | -17,61 |
| | LEU 81 | -27,07 | -17,27 | -13,65 |
| | ASN 82 | -29,14 | -32,08 | -26,21 |
| | SER 83 | -27,56 | | |
| | LYS 84 | -1,93 | | |
| | GLU 85 | -46,07 | -3,06 | -92,01 |
| | LEU 86 | -4,66 | | |
| | LEU 87 | -12,01 | -13,57 | |
| | | -483,51 | -218,01 | -192,58 |

**Supplementary Table 1** Cont'd

**ZmCYS1**

| | Residue | Interaction Energy (kcal/mol) Papain | Cathepsin L | Intestain D4 |
|---|---|---|---|---|
| N-ter | ALA 1 | -17,87 | -205,22 | -198,51 |
| | GLY 2 | -27,11 | -28,36 | -19,54 |
| | MET 3 | -31,20 | -36,39 | -15,67 |
| | LEU 4 | -19,17 | -41,81 | -26,70 |
| | ALA 5 | -18,13 | -61,31 | -23,04 |
| | GLY 6 | -13,66 | -30,00 | -29,35 |
| | GLY 7 | -4,71 | -7,37 | -12,32 |
| | LYS 9 | | -180,72 | |
| | GLN 19 | | | -1,32 |
| | | -131,86 | -591,19 | -326,45 |
| L1 | THR 49 | -18,09 | -38,63 | |
| | GLN 50 | | -42,68 | |
| | VAL 51 | -9,57 | -40,04 | -10,00 |
| | VAL 52 | -12,63 | -37,29 | -20,57 |
| | ALA 53 | -20,35 | -33,56 | -27,16 |
| | GLY 54 | -12,90 | -24,34 | -19,10 |
| | THR 55 | -44,49 | -17,41 | |
| | MET 56 | -18,39 | -15,45 | |
| | TYR 57 | | -39,84 | |
| | TYR 58 | | -33,15 | |
| | | -136,43 | -322,39 | -76,83 |
| L2 | TRP 77 | -8,12 | -36,16 | -20,84 |
| | GLU 78 | -40,45 | | -19,76 |
| | LYS 79 | 2,99 | -98,89 | -88,20 |
| | PRO 80 | -19,49 | -25,71 | -12,67 |
| | TRP 81 | -42,48 | -69,54 | -13,05 |
| | GLU 82 | -73,29 | -14,29 | 8,43 |
| | ASN 83 | -19,12 | | -23,68 |
| | PHE 84 | -25,73 | -24,64 | |
| | LYS 85 | -6,10 | | |
| | GLN 86 | -22,22 | -17,45 | -32,89 |
| | GLN 88 | -22,04 | -33,74 | |
| | | -267,92 | -284,27 | -181,83 |

**Supplementary Table 2** Primary sequences of recipient cystatin SlCYS8, donor cystatins StCYS5, PpCYS and CsCYS, and SlCYS8 LRD hybrids bearing one, two or three structural elements of StCYS5, PpCYS or CsCYS. See Table 3 for SE hybrids nomenclature. The N-terminal trunk and inhibitory loops of StCYS5, PpCYS and CsCYS are shown in bold.

| Donor | Cystatin | **N-ter trunk (N)** | α-helix | 'Elbow' region | **Loop 1 (1)** | Inter-loop region | **Loop 2 (2)** | C-ter region |
|---|---|---|---|---|---|---|---|---|
| | **SlCYS8** | 1–13<br>NPGGITNVPFPNL | 14–32<br>PQFKDLARFAVQDYNKKEN | 33–43<br>AHLEFVENLNV | 44–55<br>KEQVVAGIIYYI | 56–69<br>TLVATDAGKKKIYE | 70–84<br>TKILVKGWENFKEVQ | 85–95<br>EFKLVGDATKP |
| | StCYS5 | **1–13**<br>**KLGGFTEVPFPNS** | 14–32<br>PEFQDLTRFAVHQYNKDQN | 33–43<br>AHLEFVENLNV | **44–55**<br>**KKQVVAGMLYYI** | 56–69<br>TFAATDGGKKKIYE | **70–84**<br>**TKIWVKVWENFKKVV** | 85–94<br>EFKLVGDDSA |
| | PpCYS | **1–16**<br>**MLSGGKQEVDLQNSNN** | 17–35<br>LEIDEAAKFAVAEHNDREN | 36–48<br>SLEKLTFSKVVSC | **49–60**<br>**HMQVVAGSMYYL** | 61–74<br>VIEVEEGSSIKLYE | **75–89**<br>**AKVWVKPWQNFKKLE** | 90–101<br>EFKLKDAGVTSA |
| | CsCYS | **1–18**<br>**MASDLVPGGYTPVENPQS** | 19–37<br>SRMKELAEWAVAEHNKKAG | 38–48<br>THLMFIGILTC | **49–60**<br>**ESQIVDGVNYRF** | 61–77<br>TLTAKDEKDNCEIESYM | **78–98**<br>**AVVFEQPWEHIKELVYFQKLL** | 99–103<br>LAEQ |
| StCYS5 | S-N | **1–13**<br>**KLGGFTEVPFPNS** | 14–32<br>PQFKDLARFAVQDYNKKEN | 33–43<br>AHLEFVENLNV | 44–55<br>KEQVVAGIIYYI | 56–69<br>TLVATDAGKKKIYE | 70–84<br>TKILVKGWENFKEVQ | 85–95<br>EFKLVGDATKP |
| | S-1 | 1–13<br>NPGGITNVPFPNL | 14–32<br>PQFKDLARFAVQDYNKKEN | 33–43<br>AHLEFVENLNV | **44–55**<br>**KKQVVAGMLYYI** | 56–69<br>TLVATDAGKKKIYE | 70–84<br>TKILVKGWENFKEVQ | 85–95<br>EFKLVGDATKP |
| | S-2 | 1–13<br>NPGGITNVPFPNL | 14–32<br>PQFKDLARFAVQDYNKKEN | 33–43<br>AHLEFVENLNV | 44–55<br>KEQVVAGIIYYI | 56–69<br>TLVATDAGKKKIYE | **70–84**<br>**TKIWVKVWENFKKVV** | 85–95<br>EFKLVGDATKP |
| | S-N1 | **1–13**<br>**KLGGFTEVPFPNS** | 14–32<br>PQFKDLARFAVQDYNKKEN | 33–43<br>AHLEFVENLNV | **44–55**<br>**KKQVVAGMLYYI** | 56–69<br>TLVATDAGKKKIYE | 70–84<br>TKILVKGWENFKEVQ | 85–95<br>EFKLVGDATKP |
| | S-N2 | **1–13**<br>**KLGGFTEVPFPNS** | 14–32<br>PQFKDLARFAVQDYNKKEN | 33–43<br>AHLEFVENLNV | 44–55<br>KEQVVAGIIYYI | 56–69<br>TLVATDAGKKKIYE | **70–84**<br>**TKIWVKVWENFKKVV** | 85–95<br>EFKLVGDATKP |
| | S-12 | 1–13<br>NPGGITNVPFPNL | 14–32<br>PQFKDLARFAVQDYNKKEN | 33–43<br>AHLEFVENLNV | **44–55**<br>**KKQVVAGMLYYI** | 56–69<br>TLVATDAGKKKIYE | **70–84**<br>**TKIWVKVWENFKKVV** | 85–95<br>EFKLVGDATKP |
| | S-N12 | **1–13**<br>**KLGGFTEVPFPNS** | 14–32<br>PQFKDLARFAVQDYNKKEN | 33–43<br>AHLEFVENLNV | **44–55**<br>**KKQVVAGMLYYI** | 56–69<br>TLVATDAGKKKIYE | **70–84**<br>**TKIWVKVWENFKKVV** | 85–95<br>EFKLVGDATKP |
| PpCYS | P-N | **1–16**<br>**MLSGGKQEVDLQNSNN** | 17–35<br>PQFKDLARFAVQDYNKKEN | 36–46<br>AHLEFVENLNV | 47–58<br>KEQVVAGIIYYI | 59–70<br>TLVATDAGKKKIYE | 73–87<br>TKILVKGWENFKEVQ | 88–98<br>EFKLVGDATKP |
| | P-1 | 1–13<br>NPGGITNVPFPNL | 14–32<br>PQFKDLARFAVQDYNKKEN | 33–43<br>AHLEFVENLNV | **44–55**<br>**HMQVVAGSMYYL** | 56–69<br>TLVATDAGKKKIYE | 70–84<br>TKILVKGWENFKEVQ | 85–95<br>EFKLVGDATKP |

**Supplementary Table 2** Cont'd

| Donor | Cystatin | **N-ter trunk (N)** | α-helix | 'Elbow' region | **Loop 1 (1)** | Inter-loop region | **Loop 2 (2)** | C-ter region |
|---|---|---|---|---|---|---|---|---|
| PpCYS | P-2 | 1–13<br>NPGGITNVPFPNL | 14–32<br>PQFKDLARFAVQDYNKKEN | 33–43<br>AHLEFVENLNV | 44–55<br>KEQVVAGIIYYI | 56–69<br>TLVATDAGKKKIYE | **70–84**<br>**AKVWVKPWQNFKKLE** | 85–95<br>EFKLVGDATKP |
| | P-N1 | **1–16**<br>**MLSGGKQEVDLQNSNN** | 17–35<br>PQFKDLARFAVQDYNKKEN | 36 – 46<br>AHLEFVENLNV | **47 – 58**<br>**HMQVVAGSMYYL** | 59–72<br>TLVATDAGKKKIYE | 73–87<br>TKILVKGWENFKEVQ | 89–98<br>EFKLVGDATKP |
| | P-N2 | **1–16**<br>**MLSGGKQEVDLQNSNN** | 17–35<br>PQFKDLARFAVQDYNKKEN | 36–46<br>AHLEFVENLNV | 47–58<br>KEQVVAGIIYYI | 59–72<br>TLVATDAGKKKIYE | **73–87**<br>**AKVWVKPWQNFKKLE** | 88–98<br>EFKLVGDATKP |
| | P-12 | 1–13<br>NPGGITNVPFPNL | 14–32<br>PQFKDLARFAVQDYNKKEN | 33–43<br>AHLEFVENLNV | **44–55**<br>**HMQVVAGSMYYL** | 56–69<br>TLVATDAGKKKIYE | **70–84**<br>**AKVWVKPWQNFKKLE** | 85–95<br>EFKLVGDATKP |
| | P-N12 | **1–16**<br>**MLSGGKQEVDLQNSNN** | 17–35<br>PQFKDLARFAVQDYNKKEN | 36–46<br>AHLEFVENLNV | **47–58**<br>**HMQVVAGSMYYL** | 59–72<br>TLVATDAGKKKIYE | **73–87**<br>**AKVWVKPWQNFKKLE** | 88–98<br>EFKLVGDATKP |
| CsCYS | C-N | **1–18**<br>**MASDLVPGGYTPVENPQS** | 19–37<br>PQFKDLARFAVQDYNKKEN | 38–48<br>AHLEFVENLNV | 49–60<br>KEQVVAGIIYYI | 61–74<br>TLVATDAGKKKIYE | 75–89<br>TKILVKGWENFKEVQ | 90–100<br>EFKLVGDATKP |
| | C-1 | 1–13<br>NPGGITNVPFPNL | 14–32<br>PQFKDLARFAVQDYNKKEN | 33–43<br>AHLEFVENLNV | **44–55**<br>**ESQIVDGVNYR**F | 56–69<br>TLVATDAGKKKIYE | 70–84<br>TKILVKGWENFKEVQ | 85–95<br>EFKLVGDATKP |
| | C-2 | 1–13<br>NPGGITNVPFPNL | 14–32<br>PQFKDLARFAVQDYNKKEN | 33–43<br>AHLEFVENLNV | 44–55<br>KEQVVAGIIYYI | 56–69<br>TLVATDAGKKKIYE | **70–96**<br>**AVVFEQPWEHIKELVYFQKLL** | 97–107<br>EFKLVGDATKP |
| | C-N1 | **1–18**<br>**MASDLVPGGYTPVENPQS** | 19–37<br>PQFKDLARFAVQDYNKKEN | 38–48<br>AHLEFVENLNV | **49–60**<br>**ESQIVDGVNYRF** | 61–74<br>TLVATDAGKKKIYE | 75–89<br>TKILVKGWENFKEVQ | 90–100<br>EFKLVGDATKP |
| | C-N2 | **1–18**<br>**MASDLVPGGYTPVENPQS** | 19–37<br>PQFKDLARFAVQDYNKKEN | 38–48<br>AHLEFVENLNV | 49–60<br>KEQVVAGIIYYI | 61–74<br>TLVATDAGKKKIYE | **75–101**<br>**AVVFEQPWEHIKELVYFQKLL** | 102–112<br>EFKLVGDATKP |
| | C-12 | 1–13<br>NPGGITNVPFPNL | 14–32<br>PQFKDLARFAVQDYNKKEN | 33–43<br>AHLEFVENLNV | **44–55**<br>**ESQIVDGVNYRF** | 56–69<br>TLVATDAGKKKIYE | **70–96**<br>**AVVFEQPWEHIKELVYFQK**LL | 97–108<br>EFKLVGDATKP |
| | C-N12 | **1–18**<br>**MASDLVPGGYTPVENPQS** | 19–37<br>PQFKDLARFAVQDYNKKEN | 38–48<br>AHLEFVENLNV | **49–60**<br>**ESQIVDGVNYRF** | 61–74<br>TLVATDAGKKKIYE | **75–101**<br>**AVVFEQPWEHIKELVYFQKLL** | 102–112<br>EFKLVGDATKP |
