## Supplementary figures and images for "Harnessing the functional diversity of plant cystatins to design inhibitor variants highly active against herbivorous arthropod digestive proteases"

### Supplementary Figure 2

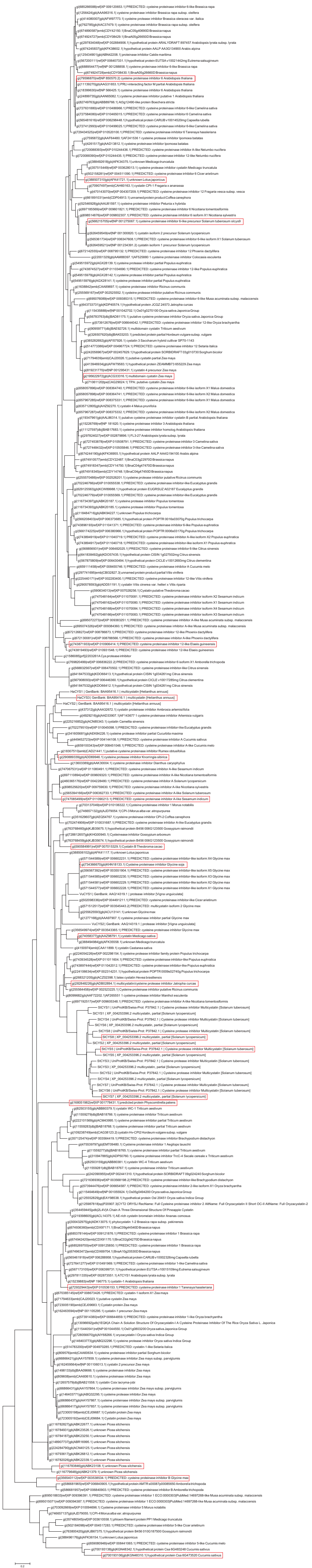
